# Autophagy promotes organelle clearance and organized cell separation of living root cap cells in *Arabidopsis thaliana*

**DOI:** 10.1101/2022.02.16.480624

**Authors:** Tatsuaki Goh, Kaoru Sakamoto, Pengfei Wang, Saki Kozono, Koki Ueno, Shunsuke Miyashima, Koichi Toyokura, Hidehiro Fukaki, Byung-Ho Kang, Keiji Nakajima

## Abstract

The root cap is a multi-layered tissue covering the tip of a plant root that directs root growth through its unique functions such as gravity-sensing and rhizosphere interaction. To prevent damages from the soil environment, cells in the root cap continuously turn over through balanced cell division and cell detachment at the inner and the outer cell layers, respectively. Upon displacement toward the outermost layer, columella cells at the central root cap domain functionally transition from gravity-sensing cells to secretory cells, but the mechanisms underlying this drastic cell fate transition are largely unknown. By using live-cell tracking microscopy, we here show that organelles in the outermost cell layer undergo dramatic rearrangements, and at least a part of this rearrangement depends on spatiotemporally regulated activation of autophagy. Notably, this root cap autophagy does not lead to immediate cell death, but rather is necessary for organized separation of living root cap cells, highlighting a previously undescribed role of developmentally regulated autophagy in plants.

**Summary statement:** Time-lapse microscope imaging revealed spatiotemporal dynamics of intracellular reorganization associated with functional transition and cell separation in the Arabidopsis root cap and the roles of autophagy in this process.

## Introduction

The root cap is a cap-like tissue covering the tip of a plant root. The root cap protects the root meristem where rapid cell division takes place to promote root elongation (Arnaud et al., 2010; Kumpf and Nowack, 2015). The root cap is also responsible for a number of physiological functions, such as gravity-sensing to redirect the root growth axis (Strohm et al., 2012), and metabolite secretion for lubrication and rhizosphere interaction (Cannesan et al., 2012; Driouich et al., 2013; Hawes et al., 2016; Maeda et al., 2019). In addition to its unique functions, the root cap exhibits a striking developmental feature, namely continuous turnover of its constituent cells (Fig. 1A) (Kamiya et al., 2016). This cell turnover is enabled by concerted production and detachment of cells at the inner stem cells layer and the outer mature cell layer, respectively. Notably, the outermost root cap cells detach from the root tip and disperse into the rhizosphere, creating a unique environment at the border between the root and the soil. For this, detaching root cap cells are called “border cells” (Hawes and Lin, 1990). Cell turnover is commonly seen in animals but rarely found in plants where morphogenesis relies not only on the production of new cells but also on the accumulation of mature and sometimes dead cells. Thus, the root cap serves as a unique experimental material to study how plant cells dynamically change their morphology and functions during tissue maintenance.

**Fig. 1.**
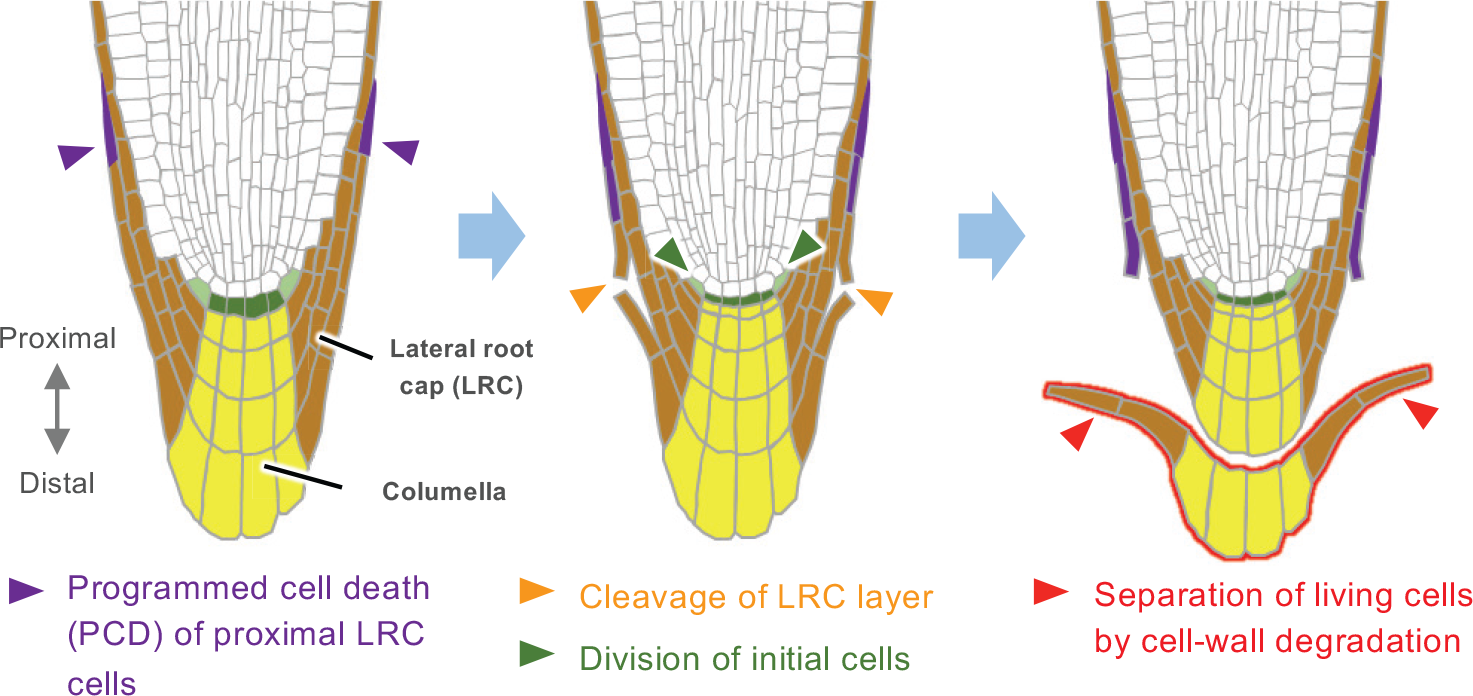
A diagram illustrating structure and cell detachment process of Arabidopsis root cap. Landmark events constituting the cell separation sequence are marked by arrowheads. Definition of the proximodistal polarity used in this study is shown on the left.

In the model angiosperm *Arabidopsis thaliana* (Arabidopsis), the root cap is composed of two radially organized domains, the central columella and the surrounding lateral root cap (LRC) that together constitute five to six cell layers along the root proximodistal axis (Fig. 1) (Dolan et al., 1993). In Arabidopsis, the outermost root cap cells do not detach individually, but rather separate as a cell layer (Fig. 1) (Driouich et al., 2007; Kamiya et al., 2016; Vicre et al., 2005). Previous studies revealed that detachment of the Arabidopsis root cap cells is initiated by localized activation of programmed cell death (PCD) at the proximal LRC region, and requires the functions of the NAC-type transcription factor SOMBRERO (SMB), a master regulator of root cap cell maturation (Bennett et al., 2010; Fendrych et al., 2014; Willemsen et al., 2008; Xuan et al., 2016). While SMB is expressed in all root cap cells and acts as a master regulator of cell maturation in the root cap, two related NAC-type transcription factors, BEARSKIN1 (BRN1) and BRN2, are specifically expressed in the outer two cell layers of the root cap (Bennett et al., 2010; Kamiya et al., 2016). BRN1 and BRN2 share high sequence similarities and redundantly promote the separation of central columella cells. Cell separation in plants requires partial degradation of cell walls. Indeed, *ROOT CAP POLYGLACTUROSE* (*RCPG*) gene encoding a putative pectin-degrading enzymes acts downstream of *BRN1* and *BRN2*, and at least BRN1 can directly bind to the *RCPG* promoter (Kamiya et al., 2016). *CELLULASE5* (*CEL5*) gene encoding a putative cellulose-degrading enzyme is also implicated in cell separation in the root cap (Bennett et al., 2010; del Campillo et al., 2004).

Previous electron microscopic studies reported profound differences in the intracellular organization between the inner and the outer root cap cells of Arabidopsis (Maeda et al., 2019; Sack and Kiss, 1989). As expected from their gravity-sensing functions, columella cells in the inner layers accumulate large amyloplasts. Amyloplasts are specialized plastids containing starch granules and known to act as statoliths in the gravity-sensing cells (statocytes) in both roots and shoots (Gilroy and Swanson, 2014). In contrast, columella cells constituting the outermost root cap layer do not contain large amyloplasts, and instead accumulate secretory vesicles (Maeda et al., 2019; Poulsen et al., 2008). Thus, the observed difference in subcellular structures correlates well with the functional transition of columella cells from gravity-sensing cells to the secretory cells (Blancaflor et al., 1998; Maeda et al., 2019; Vicre et al., 2005). Before detachment, the outermost root cap cells contain a large central vacuole, likely for the storage of various metabolites (Baetz and Martinoia, 2014). In addition, a novel role of cell death promotion has been proposed for the large central vacuole in the LRC cells (Fendrych et al., 2014).

In eukaryotes, dispensable or damaged proteins and organelles are degraded by a self-digestion process called autophagy (Mizushima and Komatsu, 2011). Autophagy initiates with expansion of isolated membranes, which subsequently form spherical structures called the autophagosomes and engulf target components. In later steps, autophagosomes fuse with vacuoles, and the content of autophagosomes degraded by hydrolytic enzymes stored in the vacuole. When eukaryotic cells are subjected to stress conditions such as nutrient starvation, autophagy is activated to recycle nutrients and maintain intracellular environments in order to sustain the life of cells and/or individuals (Mizushima and Komatsu, 2011). Autophagy plays an important role not only in stress response but also in development and differentiation, as autophagy-deficient mutants are lethal in a variety of model organisms including yeast, nematode, fruit fly, and mouse (Mizushima and Levine, 2010). Genes encoding central components of autophagy, the core *ATG* genes, are conserved in the Arabidopsis genome (Hanaoka et al., 2002; Liu and Bassham, 2012). However, under normal growth conditions, autophagy-deficient Arabidopsis mutants grow normally except for early senescence (Hanaoka et al., 2002; Yoshimoto et al., 2009). Thus roles of autophagy in plant growth and development remain largely unknown.

In this study, we revealed morphological and temporal dynamics of intracellular rearrangement that enable the functional transition of the root cap cells in Arabidopsis by using motion-tracking time-lapse imaging. We also found that the autophagy-deficient Arabidopsis mutants are defective in cell clearance and vacuolization of the outermost root cap cells. Unexpectedly, the autophagy-deficient mutants are impaired in the organized separation of the outermost root cap layer. Thus our study revealed a novel role of developmentally regulated autophagy in the root cap differentiation and functions.

## Results

### Outermost columella cells undergo rapid organelle rearrangement before cell detachment

While previous electron microscopic studies have revealed profound differences in intracellular structures between the inner and the outer root cap cells (Maeda et al., 2019; Poulsen et al., 2008; Sack and Kiss, 1989), spatiotemporal dynamics of subcellular reorganization in the root cap cells has not been analyzed, due to a difficulty in performing prolonged time-lapse imaging of the root tip that quickly relocates as the root elongates. To overcome this problem, we developed a motion-tracking microscope system with a horizontal optical axis and a spinning disc confocal unit. A similar system has been reported by another group (von Wangenheim et al., 2017). Our microscope system enabled high-magnification time-lapse confocal imaging of the tip of vertically growing roots for up to six days, allowing visualization of cellular and subcellular dynamics of root cap cells during three consecutive detachment events (Supplementary Fig. S1).

Under our experimental conditions, the outermost root cap layer of wild-type Arabidopsis sloughed off with a largely fixed interval of about 38 hours (h) (Supplementary Fig. S1F). This periodicity is comparable to that reported for roots growing on agar plates (Shi et al., 2018), indicating that our microscope system does not affect the cell turnover rate of the root cap. Bright-field observation revealed that the cell detachment initiates in the proximal LRC region and extends toward the central columella region (Fig. 1 and Fig. S1A-S1D). In concert with the periodic detachment of the outermost layer, subcellular structures of the neighboring inner cell layer (hereafter called the second outermost layer) rearranged dynamically (Fig. 2A and Supplementary Movie S1). Before the detachment of the outermost layer, columella cells in the inner three to four cell layers contained large amyloplasts that sedimented toward the distal (bottom) side of the cell (Fig. 2A, -4 h, light blue arrowheads), whereas those in the outermost layer were localized in the middle region of the cell (Fig. 2A, -4 h, dark blue arrowhead). A few hours after the outermost layer started to detach at the proximal LRC region, the amyloplasts in the second outermost layer relocated toward the middle region of the cell, resulting in a similar localization pattern to those of the outermost layer (Fig. 2A, 0.5 h, dark blue arrowheads). Toward the completion of the cell separation, rapid vacuolization and shrinkage of amyloplasts took place in the outermost layer (Fig. 2A, 18 h, green arrowhead).

**Fig. 2.**
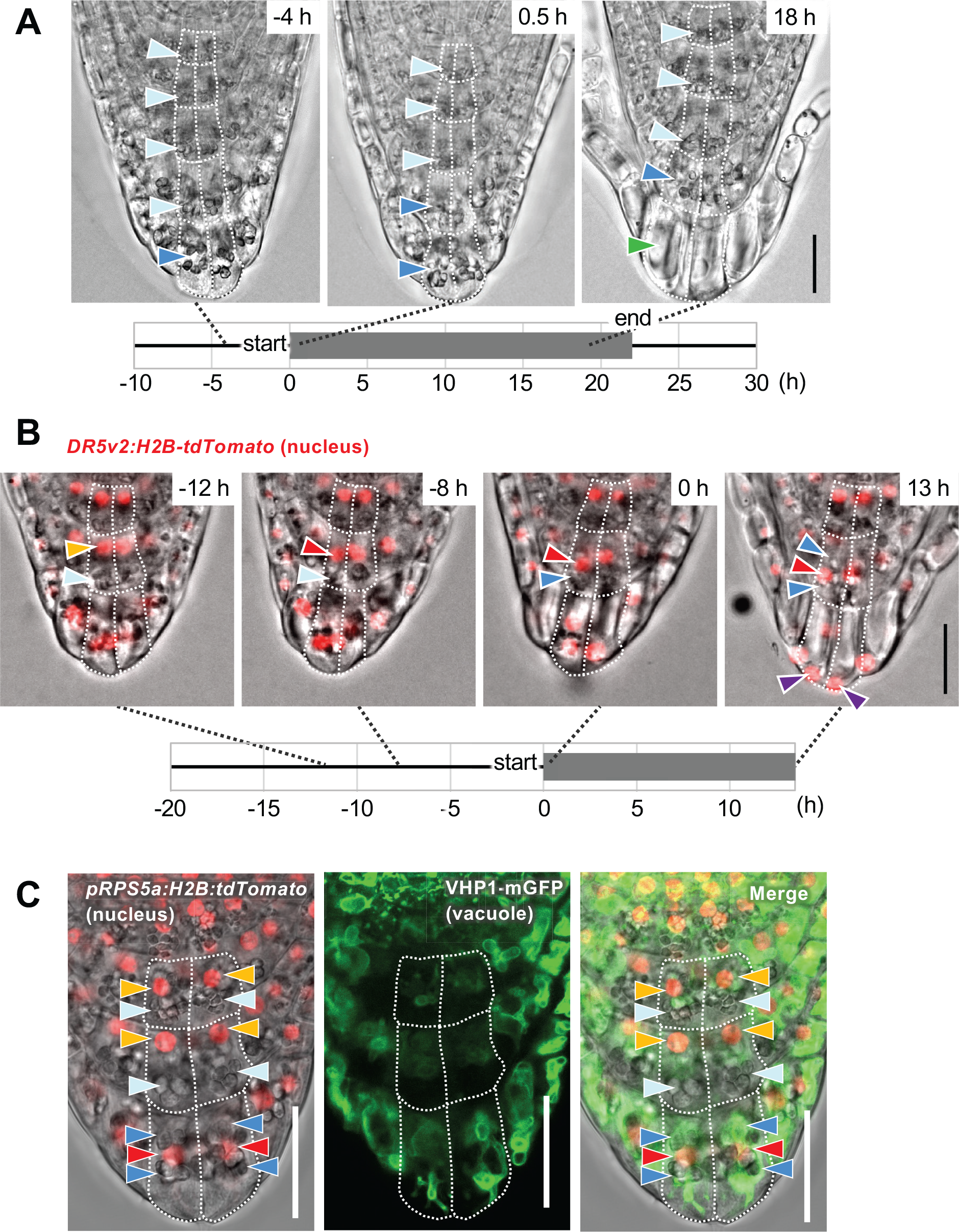
Organelle rearrangement takes place in the outer root cap layers **(A)** Time-lapse images visualizing the sequences of root cap cell detachment and relocation of amyloplasts. Representative images before (left panel), at the beginning (central panel), and around the end (right panel) of cell layer detachment are shown. Light blue and dark blue arrowheads indicate sedimenting and floating amyloplasts, respectively. Green arrowhead points to a highly vacuolated cell. Corresponding video is available as Supplementary Movie S1. **(B)** Time-lapse images showing intracellular relocation of nuclei (red fluorescence of *DR5v2:H2B-tdTomato*) and amyloplasts (gray particles in the bright field). Orange and red arrowheads point to the nuclei localized in the proximal (upper) and the middle regions of the cell, respectively. Light blue and dark blue arrowheads point to the amyloplasts in the distal (bottom) and the middle regions of the cell, respectively. Purple arrowheads point to the nuclei localized at the distal pole of the cells. Corresponding video is available as Supplementary Movie S2. **(C)** Confocal images visualizing differential localization of organelles between the inner and the outermost cell layers. Orange and red arrowheads point to red-fluorescent nuclei in the proximal (upper) and the middle regions in the cell, respectively. Light blue and dark blue arrowheads point to the amyloplasts in the distal (bottom) and the middle regions in the cell, respectively. Green color indicates vacuolar membranes. Time tables shown in (A) and (B) represent durations of the cell detachment process (gray box). Timing of image capturing is indicated at the upper right corner of each image where the origin (0 h) is set at the time when the outermost layer started detachment in the proximal LRC region. Cell outlines are delineated by white dotted lines. Scale bar, 20 µm.

By using plants expressing nuclear-localized red fluorescent proteins (*DR5v2:H2B-tdTomato*), we could also visualize dynamic relocation of nuclei, as well as its temporal relationship with amyloplast movement (Fig. 2B and Supplementary Movie S2). In the second outermost layer, nuclei relocated from the proximal (upper) to the middle region of each cell about a few hours before the neighboring outermost layer initiated detachment (Fig. 2B, -8 h, red arrowhead). This nuclear migration was followed by the relocation of amyloplasts around the time when the neighboring outermost layer initiated detachment at the proximal LRC region (Fig. 2B, 0 h, dark blue arrowhead). In later stages, the amyloplasts surrounded the centrally-localized nucleus (Fig. 2B, 13 h, dark blue arrowhead). In the outermost cells, nuclei migrated further to localize to the distal pole of the cell (Fig. 2B, 13 h, purple arrowheads).

Dynamic change in vacuolar morphology was also visualized using plants expressing a tonoplast marker (*VHP1-mGFP*) (Segami et al., 2014) (Supplementary Fig. S2 and Supplementary Movie S3). Vacuoles in the inner columella cells were smaller and spherical, whereas those in the outer cells were larger and tubular (Supplementary Fig. S2, 5-23 h). Notably, in the outermost layer, vacuoles were dramatically enlarged, and eventually occupied the entire volume of detaching root cap cells (Supplementary Fig. S2, 35-47 h). Confocal imaging of plants expressing both tonoplast and nuclear markers (*VHP1-mGFP* and *pRPS5a:H2B-tdTomato*) (Adachi et al., 2011; Segami et al., 2014) revealed that both nuclei and amyloplasts were embedded in the meshwork of vacuolar membranes in the outermost cell layer, whereas, in the inner cell layer, amyloplasts were localized in a space devoid of vacuolar membranes (Fig. 2C). Taken together, our time-lapse microscopic imaging revealed a highly organized sequence of organelle rearrangement in the outer root cap cells, as well as its close association with cell position and cell detachment.

### Autophagy is activated in the outermost root cap cells before their detachment

Autophagy is an evolutionarily conserved self-digestion system in eukaryotes and operates by transporting cytosolic components and organelles to the vacuole for nutrient recycling and homeostatic control (Mizushima and Komatsu, 2011). The rapid disappearance of amyloplasts and the formation of large vacuoles observed in the outermost root cap cells made us hypothesize that autophagy operates behind their dynamic subcellular rearrangements before the cell detachment. To test this hypothesis, we examined whether autophagosomes, spherical membrane structures characteristics of autophagy, are formed in the root cap cells at the time and space corresponding to the organelle rearrangement.

We first observed an autophagosome marker, *35Spro:GFP-ATG8a*, which ubiquitously expresses GFP-tagged Arabidopsis ATG8a proteins, one of the nine ATG8 proteins encoded in the Arabidopsis genome (Yoshimoto et al., 2004). ATG8 is a ubiquitin-like protein, and upon autophagy activation, incorporated into the autophagosome membranes as a conjugate with phosphatidylethanolamine (Liu and Bassham, 2012). Our time-lapse confocal imaging revealed uniform localization of GFP- ATG8a fluorescence in the inner cell layers, suggesting low autophagic activity in these cells (Fig. 3B and Supplementary Movie S4). In contrast, in detaching outermost cells, dot-like signals of GFP-ATG8a became evident and their number and size increased (Fig. 3C, -24.0-1.5 h). In later stages, GFP-ATG8a signals largely disappeared in the outermost cells before their detachment (Fig. 3C, 10 h). After the detachment of the outermost cell layer, the inner cells (the new outermost cells) remained showing uniform GFP-ATG8 signals (Fig. 3C, 18.5 h). In the later phase of cell detachment, GFP-ATG8a signals exhibited ring-like shapes, a typical image of autophagosomes in confocal microscopy (Fig. 3C, 1.5 h, red arrowhead and a magnified image in the inset).

**Fig. 3.**
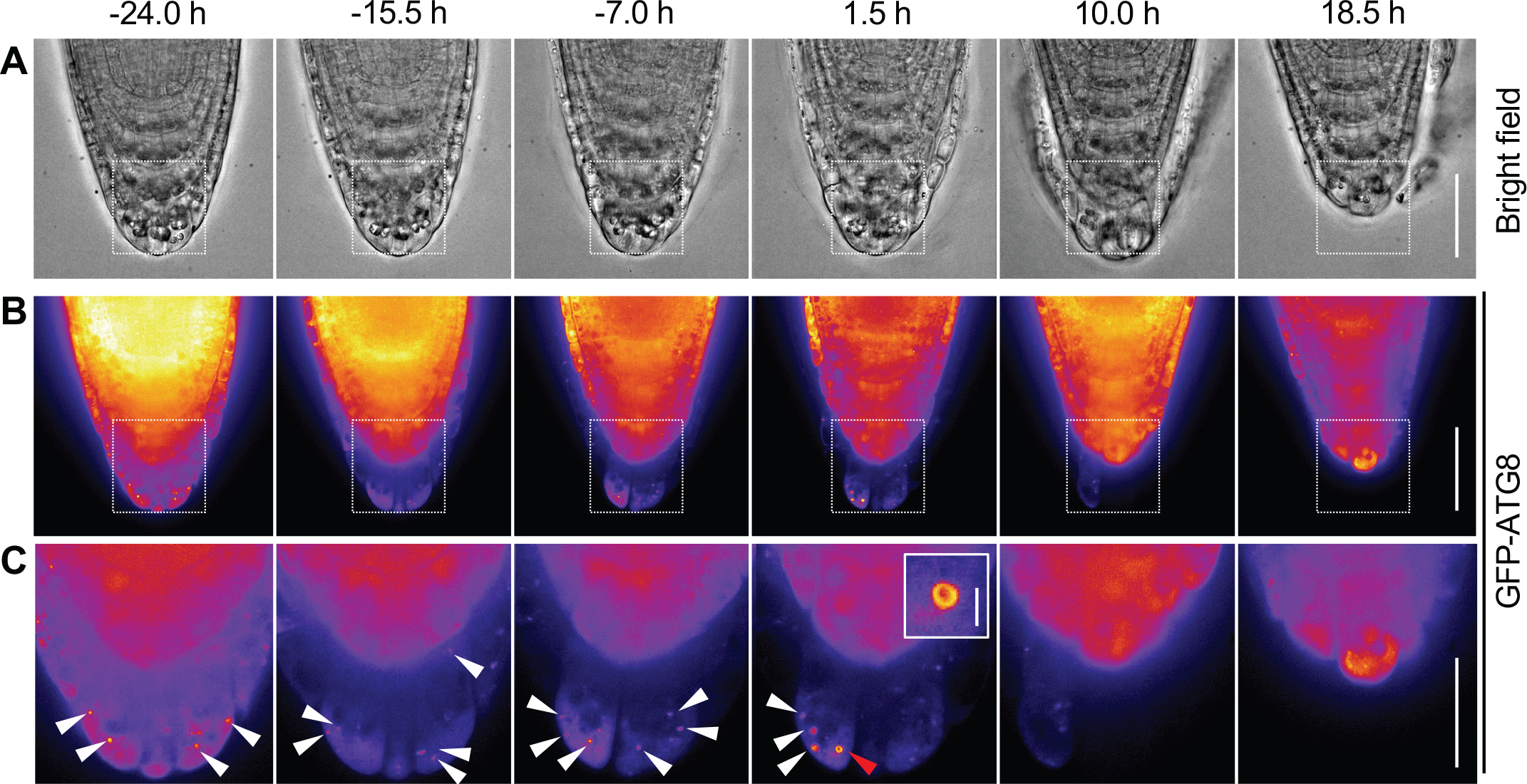
Autophagosomes are formed specifically in the outermost root cap layer Representative confocal time-lapse images of the *35Spro:GFP-ATG8a* root. Bright-field (A) and GFP-ATG8a fluorescence (B, C) images are shown. Images in (C) are magnified images of the boxed regions in (B). White arrowheads in (C) indicate autophagosomes marked by GFP-ATG8a. They showed the typical donut-shaped autophagosome images in the later phase of detachment (red arrowhead at 1.5h, inset: enlarged view). Timing of image capturing is indicated at the upper right corner of each image where the origin (0 h) is set at the time when the outermost layer started detachment in the proximal LRC region. Scale bar, 50 µm (A, B), 20 µm (C), 2 µm (C, inset). A corresponding video is available as Supplementary Movie S4.

To further confirm whether the GFP-ATG8a-labelled puncta correspond to the typical double membrane-bound autophagosome, we performed correlative light and electron microscopy (CLEM) analysis (Fig. 4) (Wang and Kang, 2020). GFP fluorescence precisely colocalized with spherical structures typical of autophagosomes (Fig. 4C-4F). Together, our observations confirmed that autophagy is activated in the outermost columella cells before their detachment.

**Fig. 4.**
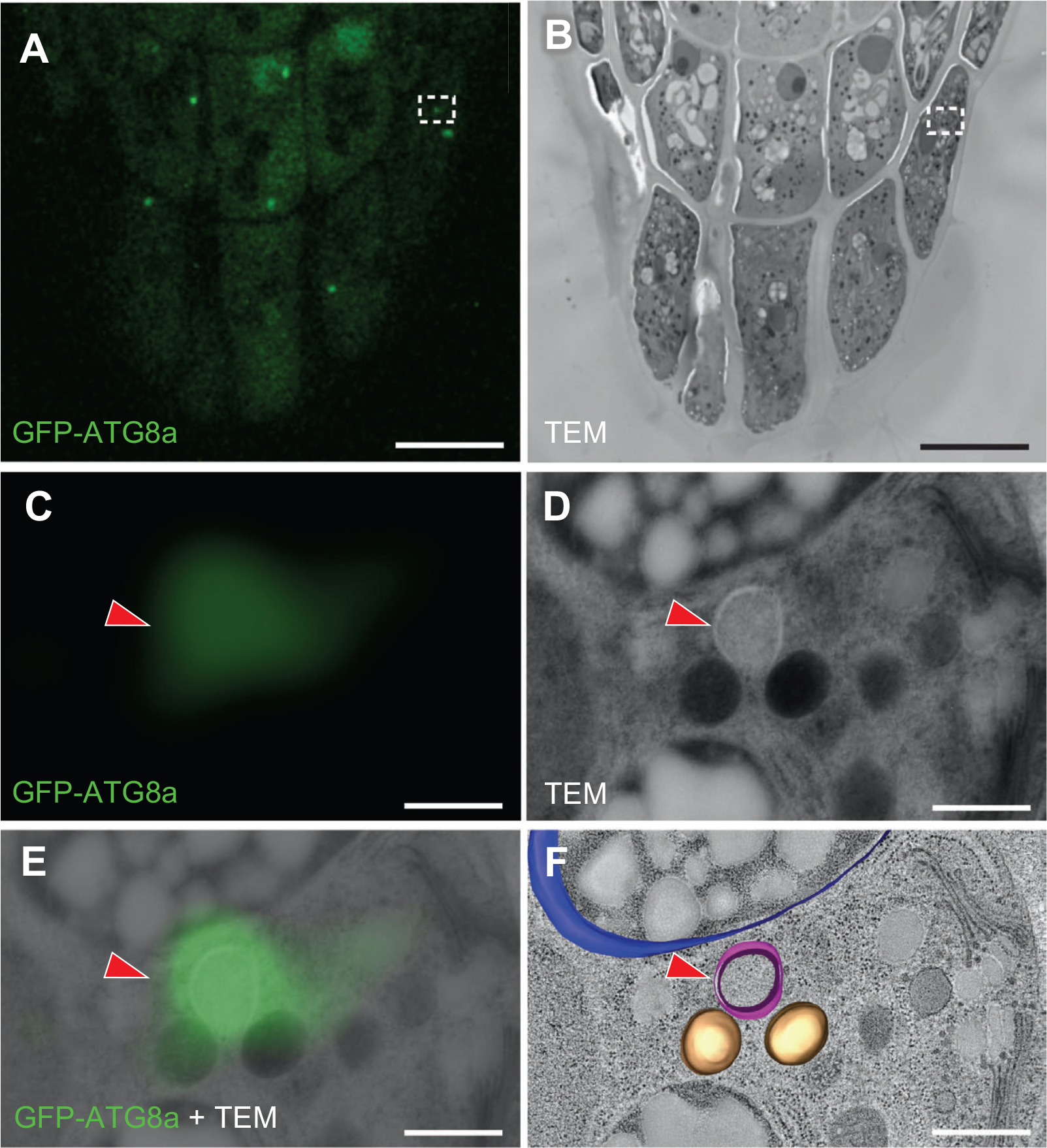
CLEM imaging revealed localization of GFP-ATG8a in autophagosomes **(A, B)** GFP fluorescence (A) and TEM (B) images of a section from a *GFP-ATG8a* root cap. **(C-E)** Magnification of the region boxed in (A) and (B). GFP-ATG8a (C), TEM (D), and their merged image (E) are shown. Red arrowhead in (E) indicates an autophagosome with GFP-ATG8a fluorescence. **(F)** A 3D electron tomographic model built for an amyloplast (blue), two mitochondria (brown,) and an autophagic compartment (magenta) overlaid with the TEM image. Scale bar, 10 µm (A, B); 500 nm (C-F).

### Autophagy promotes organelle rearrangement in the outermost root cap cells

To examine whether autophagy plays a role in the maturation of columella cells, we first tested the effect of E-64d, a membrane-permeable protease inhibitor that promotes the accumulation of autophagic bodies inside the vacuole (Inoue et al., 2006; Merkulova et al., 2014). In the outermost columella cells of E64d-treated roots, autophagic body-like aggregates accumulated inside the enlarged vacuoles, suggesting the occurrence of active autophagic degradation in these cells (Fig. S3B, compare with S3A).

We next carried out the phenotypic characterization of autophagy-deficient mutants. *ATG* genes encoding autophagy components are known to exist in the genomes of Arabidopsis and other model plant species (Hanaoka et al., 2002; Liu and Bassham, 2012). Among them, *ATG5* belongs to the core *ATG* genes and is essential for autophagosome formation as *ATG8*. In the loss of function *atg5-1* mutant (Yoshimoto et al., 2009), GFP-ATG8a signal was uniformly distributed throughout the cytosol both during and after the cell detachment, indicating that autophagosome formation in the detaching columella cells requires functional *ATG5* (Fig. S4 and Supplementary Movie S5). Furthermore, time-lapse observation revealed a loss of full vacuolation in the detaching outermost cells of *atg5-1* (Fig. S5A, Supplementary Movie S6). In the detaching outermost cells of wild-type plants, a central vacuole enlarged to occupy the entire cell volume, whereas only a few spherical and small fragmented vacuoles were found in the corresponding cells of *atg5-1* (Fig. 5A-5D). Whereas the disappearance of iodine-stained large amyloplasts was not affected in the outer columella cells of *atg5-1* (Fig. S3C and S3D), plastids in the *atg5-1* mutant exhibited abnormal morphologies dominated by tubular structures called stromules (Hanson and Hines, 2018), suggesting a specific role of autophagy in plastid restructuring and/or degradation (Fig. S3E and S3F). We also found that the detaching *atg5-1* cells were strongly stained with FDA, a compound that emits green fluorescence when hydrolyzed in the cytosol, as compared with the restricted fluorescence in the cortical region of corresponding wild-type cells (Fig. 5E and 5F). Retention of cytosol in detaching columella cells was also observed in FDA-stained roots of additional *atg* mutants including *atg2-1*, *atg7-2*, *atg10-1*, *atg12ab*, *atg13ab* and *atg18a* (Fig. 5G-5L), as well as in *atg5-1* plants expressing GUS-GFP fusion proteins under the outer layer-specific *BRN1* promoter (Fig. S5D, compare with S5C). Defects of vacuolization and cytosol digestion in *atg5-1* were complemented with an *ATG5-GFP* transgene, where GFP-tagged GFP5 proteins were expressed under the *ATG5* promoter (Fig. 5M and 5N). Together, these observations clearly demonstrated a central role of autophagy in cytosol digestion and vacuolization of detaching columella cells.

**Fig. 5.**
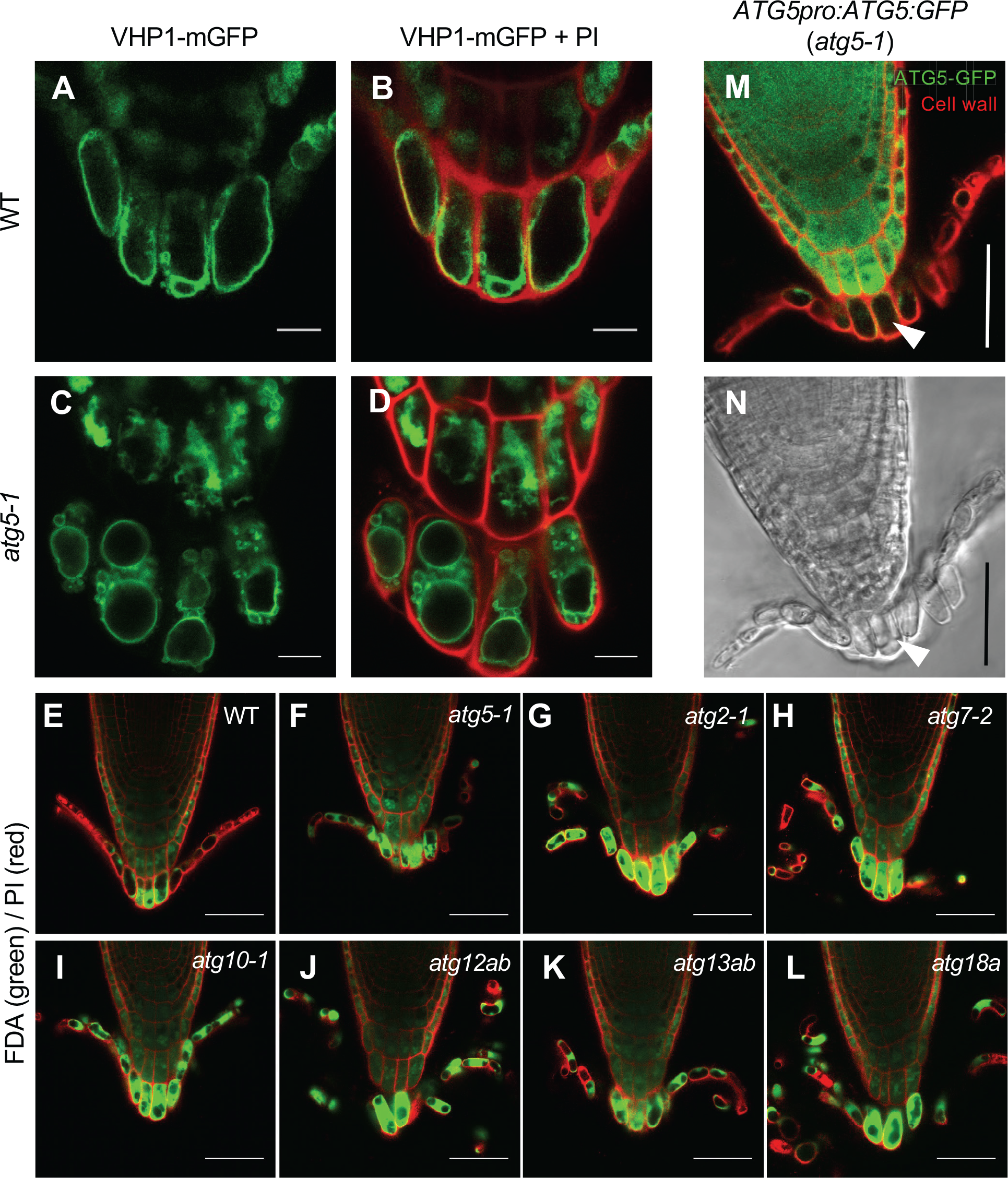
Vacuolization and cytosol digestion were inhibited in detaching columella cells in *atg* mutants **(A-D)** Vacuolar morphologies in wild-type (A, B) and *atg5-1* (C, D) columella cells. (A, C) VHP1-mGFP fluorescence (green). (B, D) Merged images with PI-stained cell walls (red). **(E-L)** Retention of cytosol in the detaching root cap cells of various *atg* mutants (F-L) as compared with wild type (E). Cytosol and cell walls were stained with FDA (green) and PI (red), respectively. **(M, N)** Vacuolization and cytosol digestion defects of detaching *atg5-1* root cap cells were complemented by the *ATG5-GFP* transgene (white arrowheads). Note the uniform ATG5:GFP expression by the *ATG5* promoter. Scale bar, 10 µm (A-D); 50 µm (E-N).

### Autophagy is required for organized separation of root cap cell layer

In the course of time-lapse imaging of *atg5-1*, we noticed that the autophagy-deficient mutants exhibited a distinct cell detachment behavior as compared with that of wild type. While the outermost root cap cells detach as a cell layer in the wild type (Fig. 6A, white arrowheads, and Supplementary Movie S7) (Kamiya et al., 2016), those of *atg5-1* detached individually (Fig. 6B, orange arrowheads, and Supplementary Movie S8), indicating that autophagy is required not only for organelle rearrangement but also for the organized separation of root cap cell layers, a behavior typically observed in the root cap of Arabidopsis and related species (Hamamoto et al., 2006; Hawes et al., 2002). The aberrant cell detachment behavior of *atg5-1* was complemented by the *ATG5-GFP* transgene (Fig. 6C, white arrowheads, and Supplementary Movie S9), confirming the causal relationship. To clarify whether autophagy activation in the outermost cells is sufficient for organized cell separation, we established *atg5-1* plants expressing GFP-tagged ATG5 proteins under the *BRN1* and the *RCPG* promoter, which drive transcription in the outer two cell layers and the outermost root cap layer, respectively (Kamiya et al., 2016). Time-lapse imaging revealed that both of the plant lines restored the organized separation of the outermost root cap cell layer (Fig. 7A and 7B, white arrowheads and Supplementary Movie S10 and S11). These observations, in particular, restoration of the layered cell separation by the *RCPG* promoter-driven ATG-GFP, confirmed that autophagy activation in the detaching cells at the timing of active cell wall degradation is sufficient for the organized separation of the outermost root cap layer.

**Fig. 6.**
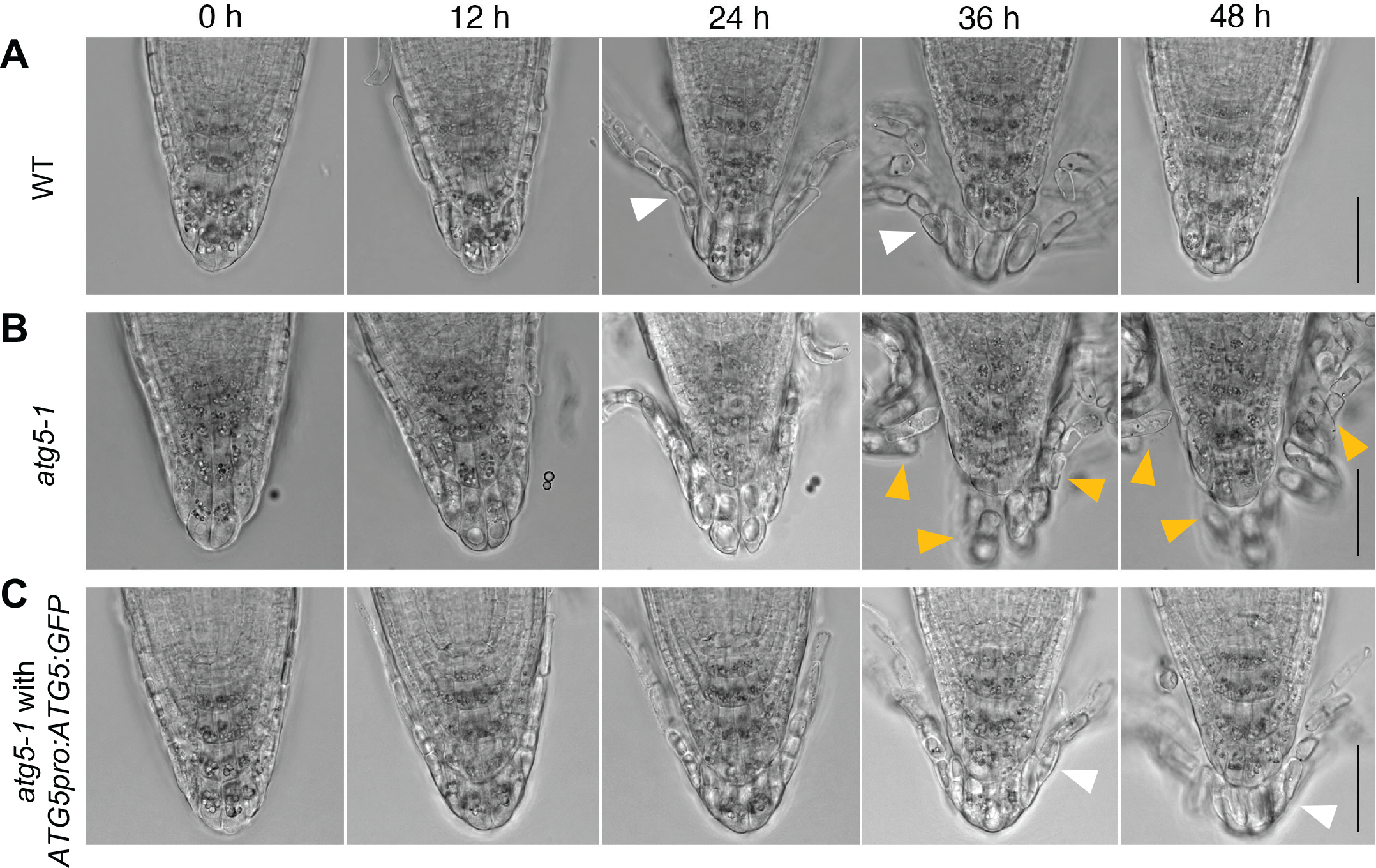
Autophagy activation is required for organized separation of the outermost root cap cell layer (A-C) Time-lapse images of root cap detachment processes in wild-type (A), *atg5-1* (B), and *ATG5pro:ATG5:GFP atg5-1* (C) plants at the time points indicated at the top. Note that the outermost root cap cells detach as a layer (white arrowheads) in wild type (A) and *ATG5:GFP atg5-1* (C), whereas they detach individually in *atg5-1* (B, orange arrowheads). Scale bar, 50 µm. Corresponding videos are available as Supplementary Movie S7-S9.

**Fig. 7.**
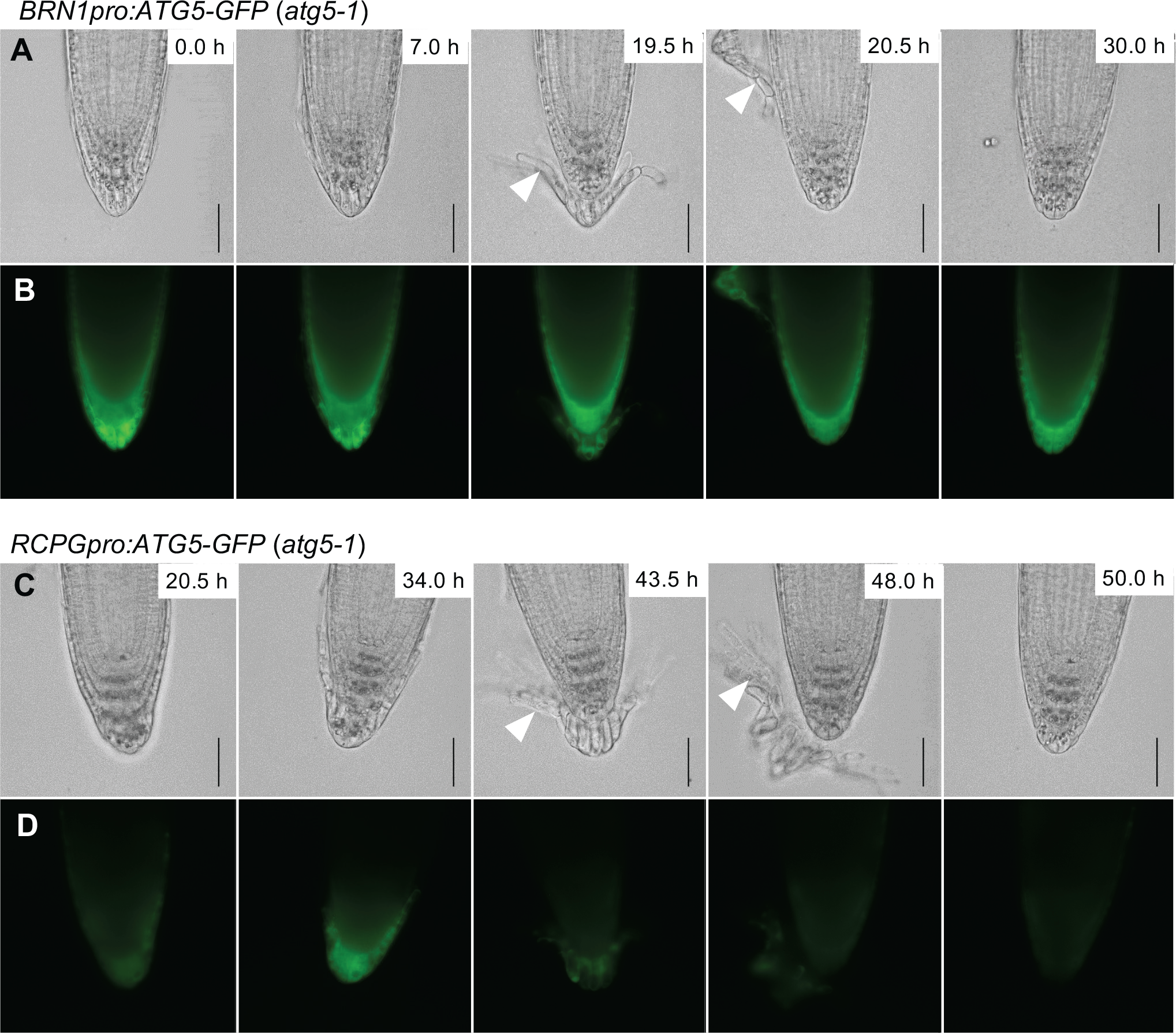
Autophagy activation at the timing of cell wall degradation is sufficient for organized cell separation **(A-D)** Time-lapse images of root cap detachment processes in *BRN1pro:ATG5-GFP atg5-1* (A, B) and *RCPGpro:ATG5:GFP atg5-1* (C, D) at the time points indicated at the top right corner of each panel. Note that the outermost root cap cells detach as a cell layer in both genotypes (white arrowheads), as compared with individual detachment in *atg5- 1* (Fig. 6B). Bright-field (A, C) and GFP fluorescence (B, D) images were shown. Scale bar, 50 µm. Corresponding videos are available as Supplementary movies S10 and S11.

## Discussion

In this study, we revealed spatiotemporal dynamics of the intracellular reorganization and cell detachment in the Arabidopsis root cap, as well as a role of developmentally regulated autophagy in these processes. In the outermost root cap layer, autophagy is activated in a specific cell layer and at the timing closely associated with the functional transition of columella cells and their detachment. This spatiotemporally regulated activation of autophagy is essential not only for cell clearance and vacuolar enlargement but also for the organized separation of the outermost layer of the root cap.

### Motion-tracking time-lapse imaging revealed rapid intracellular rearrangement associated with the functional transition of root cap cells

Cells constituting the root cap constantly turn over by balanced production and detachment of cells at the innermost and the outermost cell layers, respectively. During their lifetime, columella cells undergo a functional transition from being gravity-sensing statocytes to secretory cells according to their position (Blancaflor et al., 1998; Maeda et al., 2019; Sack and Kiss, 1989; Vicre et al., 2005). While the previous electron microscopic observations revealed a profound difference in the subcellular structures between the inner statocytes and the outer secretory cells of the Arabidopsis root cap (Maeda et al., 2019; Poulsen et al., 2008; Sack and Kiss, 1989), detailed temporal dynamics of organelles rearrangement in relation to the timing of cell displacement and detachment has not been analyzed.

Our time-lapse observation using a motion-tracking microscope system with a horizontal optical axis clearly visualized both morphological and temporal details of organelle rearrangement in this transition (Fig. 8). Cells in the inner two to three layers have unique arrangements of organelles, which is likely optimized for their gravity-sensing function (Blancaflor et al., 1998). In these cells, starch granule-containing amyloplasts and nuclei are localized at the distal (lower) and proximal (upper) end of each cell, respectively, whereas small tubular vacuoles preferentially occupy the proximal (upper) half of each cell (Fig. 2) (Leitz et al., 2009; Sack and Kiss, 1989). This organelle arrangement changed dynamically in the outermost cell layer. The first conspicuous sign of rearrangement is relocation of nuclei from the upper to the central region, which happens even before the layer containing these columella cells starts to detach at the proximal LRC region (Fig. 2). Around the time of the detachment of this cell layer, amyloplasts ’float up’ to the middle region of the cell (Fig. 2). Later, amyloplasts disappear and vacuoles start to expand to occupy the entire cell volume by the time these cells slough off from the root tip (Fig. 2 and Supplementary Fig. S2). The development of large central vacuoles likely constitutes a central component of functional specialization of these cells for storage (Driouich et al., 2013; Hawes et al., 2016; Vicre et al., 2005). A novel role of central vacuoles for cell death promotion has been also proposed for LRC cells (Fendrych et al., 2014).

**Fig. 8.**
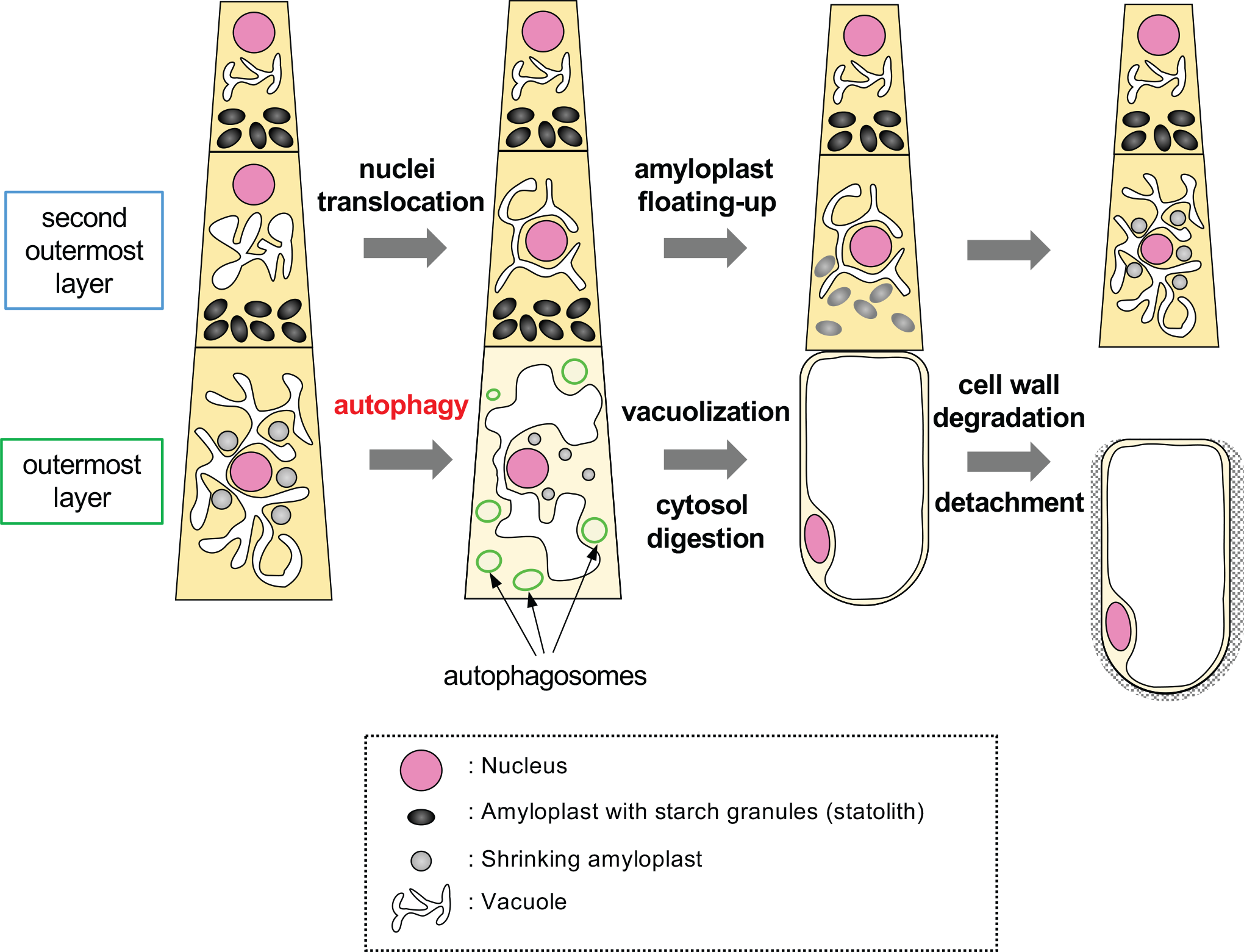
Schematic illustration of the sequence of organelle rearrangement and autophagy activation during maturation and detachment of columella cells.

Here, the central question is what controls the spatiotemporal activation of this dramatic rearrangement of organelles in the root cap. The NAC-type transcription factors BRN1 and BRN2 are expressed specifically in the outer two cell layers of the root cap and required for cell detachment (Bennett et al., 2010; Kamiya et al., 2016), seemingly becoming good candidates for the upstream regulators. However, the outermost root cap cells of *brn1 brn2* mutants, though defective in cell detachment, were found to be normally vacuolated and lacking amyloplasts as those of wild type, indicating that at least a part of the organelle rearrangement is regulated independently of *BRN1* and *BRN2* (Bennett et al., 2010; Kamiya et al., 2016). On the other hand, our previous study suggested the existence of unknown positional cues that, together with another NAC-type transcription factor SMB, promote the outer layer-specific expression of *BRN1* and *BRN2* (Kamiya et al., 2016). Future identification of factors transmitting such positional information will provide a clue to understanding a mechanism underlying position-dependent organelle rearrangement in the root cap.

### Autophagy is activated in the outermost root cap cells to promote cell clearance and vacuolization

Our time-lapse imaging revealed specific activation of autophagy in the outermost root cap layer in concert with the progression of the cell separation (Fig. 3). As expected, mutants defective in the canonical autophagy pathway exhibited compromised cell clearance and vacuolization of detaching root cap cells (Fig. 5). Because detached root cap cells are dispersed into the rhizosphere and act in plant defense through their secretory capacity (Driouich et al., 2013; Hawes et al., 2016), degradation of starch-containing amyloplasts and vacuolar expansion appear to be a reasonable differentiation trajectory in view of energy-recycling and storage.

Autophagosomes are double-membrane vesicles that engulf a wide range of intracellular components and transport them to vacuoles for degradation by lytic enzymes. Rapid reduction of GFP-ATG8a signals and accumulation of autophagic body-like structures inside the vacuoles after the application of the proteinase inhibitor E64d (Supplementary Fig. S3) support occurrence of active autophagic flow and vacuolar degradation in the outermost root cap layer. Such active autophagic transport may act to supply membrane components and to facilitate water influx into the vacuoles by increasing osmotic pressure, leading to enhanced vacuolization of the outermost root cap cells.

While the autophagy-deficient *atg5-1* mutant was capable of eliminating Lugol-stained amyloplasts from mature columella cells as the wild type, morphology of plastids in the detaching root cap cells was abnormal in *atg5-1*, having tubular structures typical of stromules (Supplementary Fig. S3). Storomules arise from chloroplasts under starvation or senescence conditions. In such stress conditions, chloroplast contents are degraded via piecemeal-type organelle autophagy, in which stromules or chloroplast protrusions are believed to be engulfed by an autophagosome (Ishida et al., 2008), whereas damaged chloroplasts can be engulfed as a whole by an isolated membrane and transported into vacuoles (Izumi et al., 2013). Stromule formation in the autophagy-deficient *atg5-1* mutant suggests that amyloplast degradation in the outermost root cap cells proceeds in two steps; first by autophagy-independent degradation of starch granules and stromule formation, followed by the piecemeal chloroplast autophagy. It should be noted, however, that autophagy-dependent amyloplast degradation also occurs as a part of root hydrotropic response, where some starch-containing amyloplasts are engulfed directly by the autophagosome-like structures (Nakayama et al., 2012). Together, these observations suggest that multiple amyloplast degradation pathways exist in the Arabidopsis root cap with different contributions of autophagy.

While the present study clearly demonstrated the role of autophagy in the organelle rearrangement in the root cap, spatiotemporal regulation of autophagy activation is yet to be investigated. The root cap autophagy seems to operate via canonical macro-autophagy pathway mediated by the components encoded by the *ATG* genes (Fig. 5) (Liu and Bassham, 2012) (Fig. 5). Autophagy is induced by various stress conditions, such as nutrient starvation, as well as abiotic and biotic stresses, where SNF-related kinase 1 (SnRK1) and target of rapamycin (TOR) protein kinase complexes function as key regulators (Liu and Bassham, 2012; Mizushima and Komatsu, 2011). In contrast, the root cap autophagy can occur in plants growing on a sterile nutrient-rich medium in our experiments, suggesting that root cap autophagy is activated independently of nutrient starvation and biotic stress. Instead, activation of the root cap autophagy appears to be closely associated with the process of cell detachment, which in turn is known to be regulated by intrinsic developmental programs (Dubreuil et al., 2018; Shi et al., 2018). Again, *BRN1* and *BRN2* are unlikely to regulate the root cap autophagy, because cell clearance and vacuolization normally occur in the outermost root cap cells of *brn1 brn2* mutants.

### Autophagy is required for the organized separation of the Arabidopsis root cap cells

Autophagy promotes organelle rearrangement associated with the differentiation of secretory cells that subsequently slough off to disperse into the rhizosphere. Based on this, we expected that the loss of autophagy would inhibit or delay cell detachment in the root cap. Somewhat unexpectedly, however, autophagy-deficient *atg5-1* mutants showed a phenotype suggestive of enhanced cell detachment (Fig. 6). In Arabidopsis and related species, the outermost root cap cells separate as a cell layer, rather than as isolated cells (Driouich et al., 2010; Driouich et al., 2007; Kamiya et al., 2016). Although the physiological significance of this detachment behavior has not been demonstrated so far, it has been hypothetically linked with a capacity of secreting mucilage, a mixture of polysaccharides implicated in plant defense, aluminum-chelating, and lubrication (Driouich et al., 2010; Maeda et al., 2019).

Previous genetic studies suggested a key role of cell wall pectins in the control of root cap cell detachment; when pectin-mediated cell-cell adhesion was compromised by mutations in genes encoding putative pectin-synthesizing enzymes or overexpression of RCPG, a root cap-specific putative pectin-hydrolyzing enzyme, root cap cells slough off as isolated cells (Driouich et al., 2010; Kamiya et al., 2016). Moreover, the morphology of detaching root cap cell layers was altered in the loss-of-function *rcpg* mutant, likely due to a failure of separating cell-cell adhesion along the lateral cell edge (Kamiya et al., 2016). The similarity between the altered cell detachment behaviors between *atg5-1* and pectin-deficient plants suggests a role of autophagy in the control of cell wall integrity during the root cap cell detachment. Both transport and modification of cell wall pectins require Golgi and Golgi-derived vesicles (Driouich et al., 2012; Wang et al., 2017). In outer root cap cells, small vesicles accumulate for their secretory functions (Driouich et al., 2013; Maeda et al., 2019; Wang et al., 2017), and a mutation disrupting this secretory pathway results in the failure of root cap cell detachment (Poulsen et al., 2008). If autophagy is required for timely attenuation of such vesicular transport during the cell detachment program, lack of autophagy should lead to prolonged secretion of cell wall modifying enzymes such as RCPG, resulting in enhanced loosening of cell-cell adhesion. Indeed, we could recognize broader gaps at the apoplastic junctions at the distal cell-cell adhesion points in *atg5-1* than those in the wild type (Supplementary Movie S7 and S8). Future studies comparing secretory dynamics of cell wall-modifying enzymes in various genetic backgrounds using our live-imaging system will elucidate the molecular mechanism controlling the cell detachment behaviors in the root cap and the role of autophagy.

In summary, our study revealed the role of spatiotemporally regulated autophagy in cell clearance and vacuolization in root cap differentiation as well as in cell detachment. While autophagy has been known to promote tracheary element differentiation in Arabidopsis and anther maturation in rice, roles of autophagy in these instances are linked to PCD (Escamez et al., 2016; Kurusu and Kuchitsu, 2017). Considering that autophagy is required for functional transition and detachment of living columella cells, our study revealed a previously undescribed role of developmentally regulated autophagy in plant development.

## Materials and Methods

### Plant materials and growth conditions

*Arabidopsis thaliana* L. Heynh (Arabidopsis) accession Col-0 was used as the wild type. The Arabidopsis T-DNA insertional lines, *atg5-1* (SAIL_129_B07), *atg7-2* (GK- 655B06), *atg2-1* (SALK_076727), *atg10-1* (SALK_084434), *atg12a* (SAIL_1287_A08), *atg12b* (SALK_003192), *atg13a* (GABI_761_A11), *atg13b* (GK-510F06) and *atg18a* (GK_651D08) have been described previously (Doelling et al., 2002; Hanaoka et al., 2002; Izumi et al., 2013; Thompson et al., 2005; Yoshimoto et al., 2004; Yoshimoto et al., 2009). *35Spro:CT-GFP*, *RPS5apro:H2B-tdTomato* and *VHP1-mGFP* has been described previously (Adachi et al., 2011; Köhler et al., 1997; Segami et al., 2014). Seeds were grown vertically on Arabidopsis nutrient solution supplemented with 1 % (w/v) sucrose and 1 % (w/v) agar under the 16h light/8h dark condition at 23 °C.

### Generation of transgenic plants

For *ATG5pro:ATG5:GFP*, a 4.5-kb genomic fragment harboring the ATG5 coding region and the 5’-flanking region was amplified by PCR and cloned into pAN19/GFP-NOSt vector, which contained GFP-coding sequence and the *Agrobacterium (Rhizobium)* nopaline synthase terminator (NOS). The resulting *ATG5- GFP* fragment was then transferred to *pBIN4* to give *ATG5pro:ATG5:GFP/pBIN41*.

Layer-specific rescue constructs of *ATG5-GFP* were constructed by amplifying the *ATG5-GFP* fragment from *ATG5pro:ATG5:GFP/pBIN41*, and inserting them to pDONR221 by the GatewayTM technology. The *ATG5-GFP* fragment was then transferred to *pGWB501:BRN1pro* and *pGWB501:RCPGpro*, which respectively contained the *BRN1* and *RCPG* promoter flanking the Gateway cassette in pGWB501 (Nakagawa et al., 2007). The cytosolic marker *GUS-GFP* was similarly constructed by inserting a *GUS-GFP* fragment into pENTR D-TOPO, and then by transferring the insert to *pGWB501:BRN1pro* to give *BRN1pro:GUS-GFP*.

For *DR5v2:H2B:tdTomato,* a *DR5v2* promoter fragment was amplified by PCR from the *DRv2n3GFP* construct (Liao et al., 2015), and inserted into pGWB501 by the In-Fusion technique to give *pGWB501:DR5v2*. The *H2B-tdTomato* fragment in pENTR was transferred to the *pGWB501:DR5v2*. Integrity of the cloned genes was verified by DNA sequencing. Transformation of Arabidopsis plants was performed by the floral dip method using *Rhizobium* (formerly *Agrobacterium*) *tumefaciens,* strain C58MP90.

### Microscopy

Time-lapse imaging of the root cap was performed using two microscopic systems developed in the corresponding authors’ laboratory, which can automatically track the tip of vertically growing roots. Technical details will be published elsewhere. Briefly, an inverted microscope (ECLIPSE Ti-E and ECLIPSE Ti2-E, Nikon, Tokyo, Japan) was tilted by 90 degrees to vertically orient the sample stage. The motorized stage was controlled by the Nikon NIS-elements software with the “keep object in view” plugin to automatically track the tip of growing roots. Three-day-old seedlings were transferred to a chamber slide (Lab-Tek chambered coverglass, Thermofisher, Waltham, MA) and covered with a block of agar medium.

Confocal laser scanning microscopy was carried out with a Nikon C2 confocal microscope. Roots were stained with 10 µg/ml of propidium iodide (PI). Fluorescein diacetate (FDA) staining was performed by soaking the roots in a solution containing 2 μg/ml of FDA.

Iodine staining was performed as described previously (Segami et al., 2018). Root fixed in 4% (w/v) paraformaldehyde in PBS for 30 min under a vacuum at room temperature. The fixed sample was washed twice for 1 min each in PBS and cleared with ClearSee (Kurihara et al., 2015). The samples were transferred to 10% (w/v) xylitol and 25% (w/v) urea to remove sodium deoxycholate, and then stained in a solution containing 2 mM iodine (Wako), 10 % (w/v) xylitol, and 25 % (w/v) urea.

Correlative light and electron microscopy (CLEM) analysis was performed as described previously (Wang and Kang, 2020; Wang et al., 2019). GFP-ATG8a seedlings were grown vertically under 16 h light-8 h dark cycle at 22 °C for seven days. Root tips samples expressing GFP were cryofixed with an EM ICE high-pressure freezer (Leica Microsystems, Austria) and embedded in Lowicryl HM20 resin at -45°C. TEM sections of 150nm thickness were collected on copper or gold slot grids coated with formvar and examined for GFP after staining the cell wall with Calcofluor White. The grids were post- stained and GFP-positive cells were imaged under an H-7650 TEM (Hitachi High-Tech, Japan) operated at 80kV. For electron tomography, tilt series were collected with a TF- 20 intermediate voltage TEM (Thermo Fisher Scientific, USA). Tomogram calculation and three-dimensional model preparation were carried out with the 3dmod software package (<u>bio3d.colorado.edu</u>).

## Acknowledgments

We thank Masanori Izumi (RIKEN, Japan), Kohki Yoshimoto (Meiji University, Japan), Masayoshi Maeshima (Nagoya University, Japan), Shoji Segami (NIBB, Japan), and Maureen R. Hanson (Cornell University, USA) for providing plant materials, Dolf Weijers (Wageningen University, Netherlands) for providing the DR5v2 construct, and Masako Kanda for technical assistance.

---

**Fig. S1.**
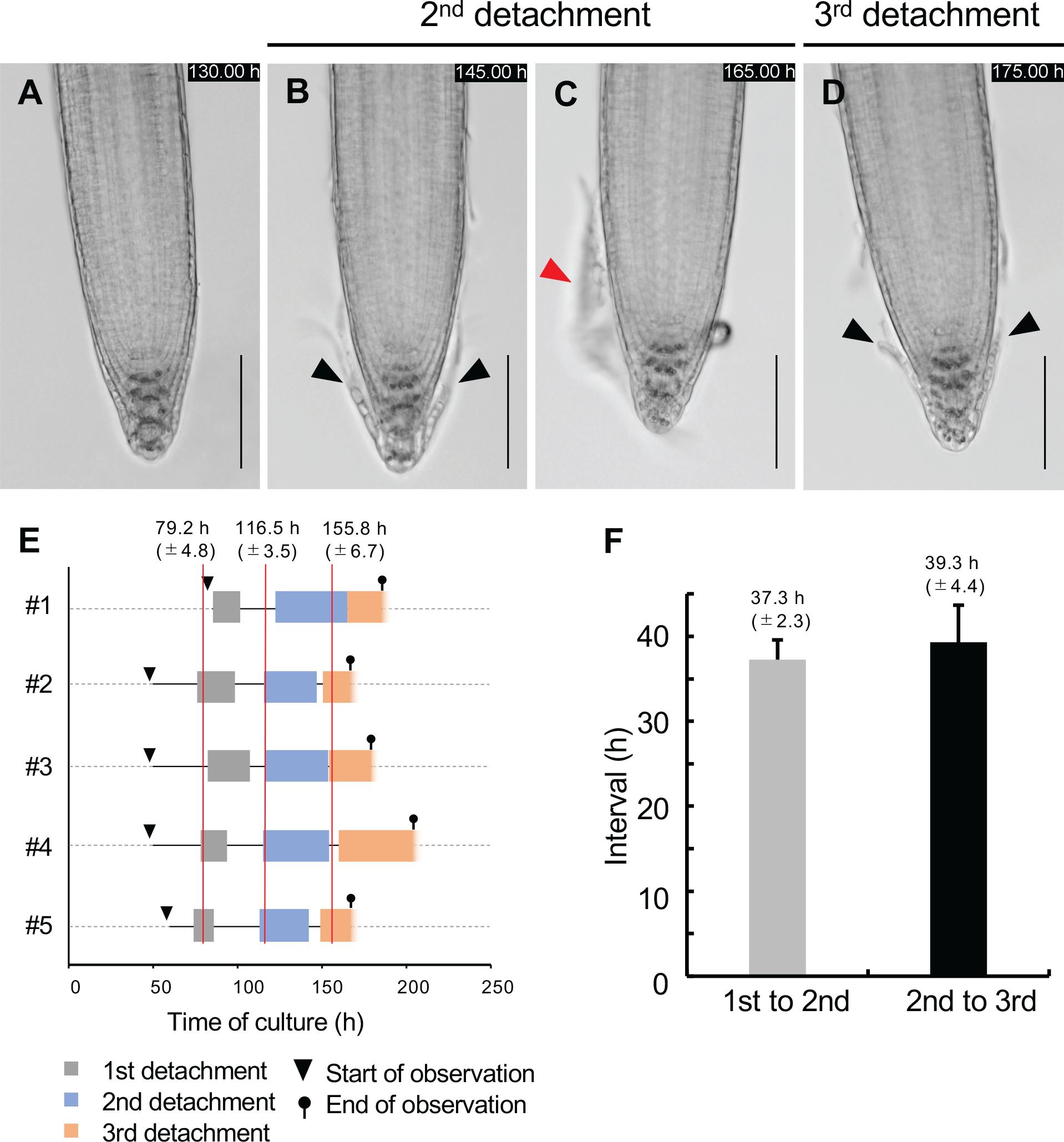
Arabidopsis root cap cells detach at fixed intervals **(A-D)** Time-lapse images showing periodic detachment of *Arabidopsis* root cap cells. Detachment of the outermost root cap layer initiates at the proximal LRC region and progressively extends toward the central columella region (B, black arrowheads). Detached root cap cells adhere together to keep a cell layer morphology (C, red arrowhead). Detachment of the next cell layer initiates in the same manner as the previous one (D). Elapsed time after the start of culture is indicated in each panel. Scale bar, 100 µm. **(E)** A time table showing periodic detachment of root cap cell layers in five (#1-5) root samples each experiencing three rounds of root cap detachment. Gray, blue, and orange boxes indicate the duration from the start (initial detachment at the proximal LRC region) and the end (complete detachment at the columella region) of the first, second, and third cell layer, respectively. The x-axis indicates elapsed time (h) from the start of culture. Red lines indicate average time points of the start of detachment. **(F)** Intervals between the start of detachment between the first and second cell layers (gray bar), and between the second and third cell layer (black bar). Mean and SE are shown (*n* = 5).

**Fig. S2.**
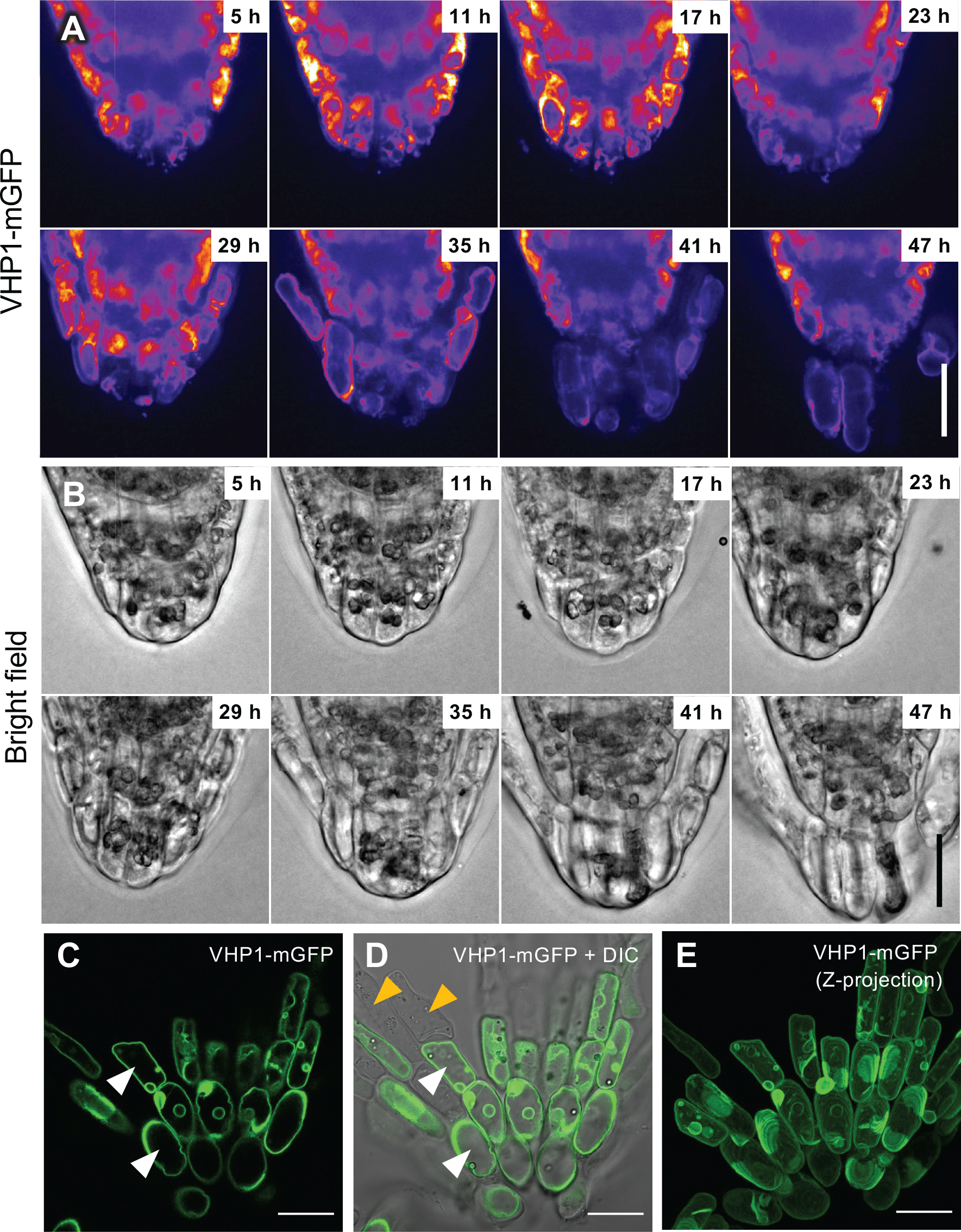
Morphological transition of vacuoles during the detachment of root cap cells **(A, B)** Time-lapse images showing vacuolar morphology by the tonoplast-localized VHP1-mGFP fluorescence (A) and bright-field images (B). In the outermost cells, vacuoles are initially small and fragmented (up to 17 h), and gradually expand to form large central vacuoles before the cell detachment (41 h). Elapsed time after the start of observation is indicated in each panel. A corresponding video is available as Supplementary Movie S3. **(C-E)** The entire cell volume was occupied by a large central vacuole in detaching root cap cells. Images of VHP1-mGFP fluorescence (C) and its overlay with a DIC image (D) were shown. (F) is a Z-stack projection encompassing 50-µm depth. Note that cells at the center of the detached cell layer possess large central vacuoles as visualized by VHP1- mGFP (white arrowheads), whereas those at the periphery do not show fluorescence (orange arrowheads) likely due to the loss of cell viability. Scale bar, 20 µm.

**Fig. S3.**
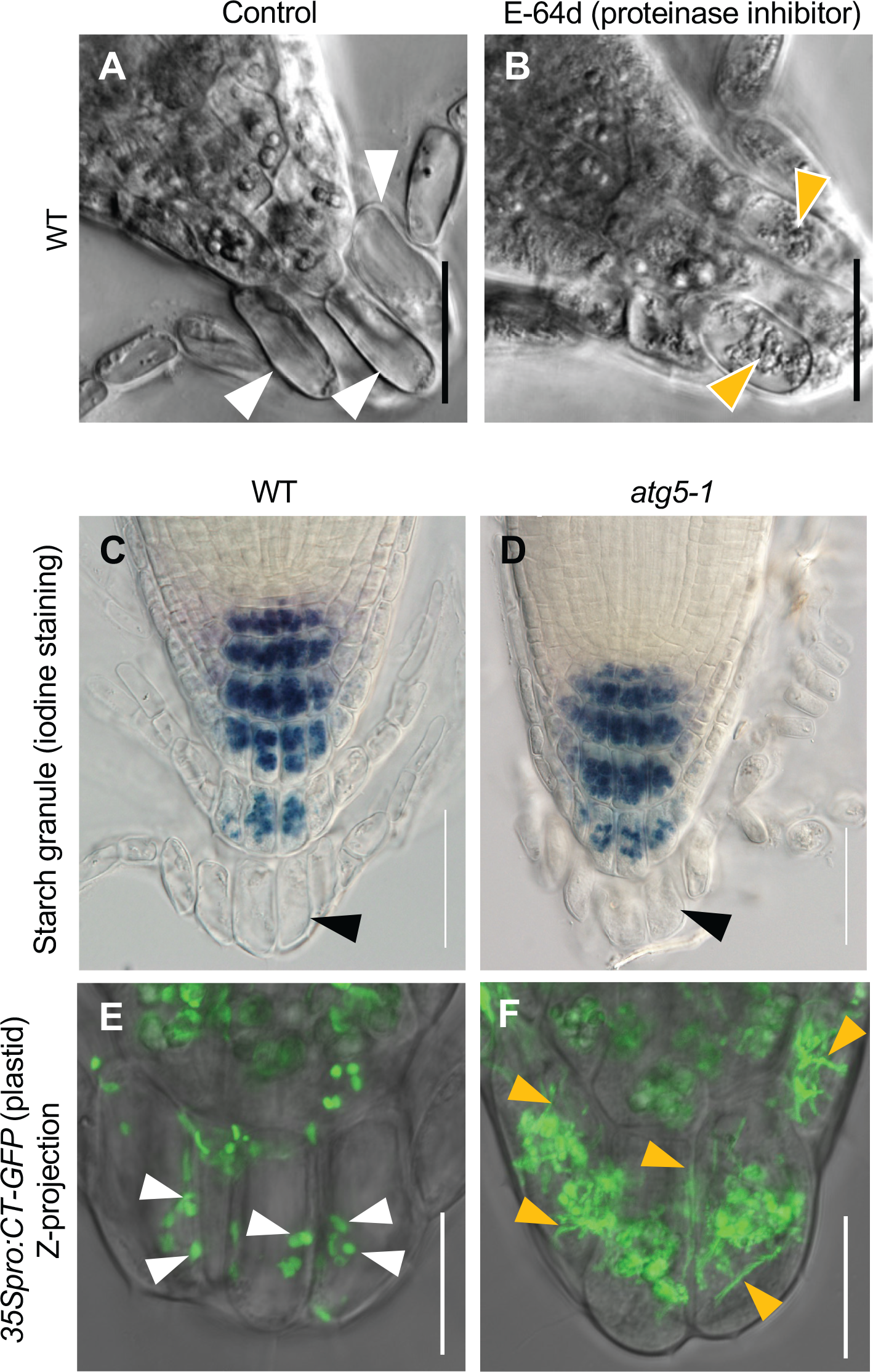
Accumulation of autophagic body-like structures in the E64d-treated wild-type root cap cells and abnormal plastid morphology in *atg5-1* **(A, B)** Accumulation of autophagic body-like structures inside the vacuoles of the wild-type outermost root cap cells after E-64d treatment (B, orange arrowheads), as compared with the translucence vacuolar images of a non-treated control (A, white arrowheads). 5- day-old seedlings grown on the medium with or without 10 µM E-64d were observed. Scale bar, 20 µm. **(C, D)** Amyloplasts in the outermost root cap cells lost starch granules in both wild type and *atg5-1*. Black arrowheads indicate the detaching outermost cell layers. Scale bar, 50 µm. **(E, F)** Amyloplasts exhibit abnormal morphologies in the outermost root cap cells of *atg5-1* (F) as compared with those in the wild type (E). Plastids are visualized by the CT-GFP fluorescence marker line. Note that small spherical plastids accumulate in the wild-type cells (white arrowheads), whereas those with tubular morphologies dominate in *atg5-1* cells (orange arrowheads). Scale bar, 20 µm.

**Fig. S4.**
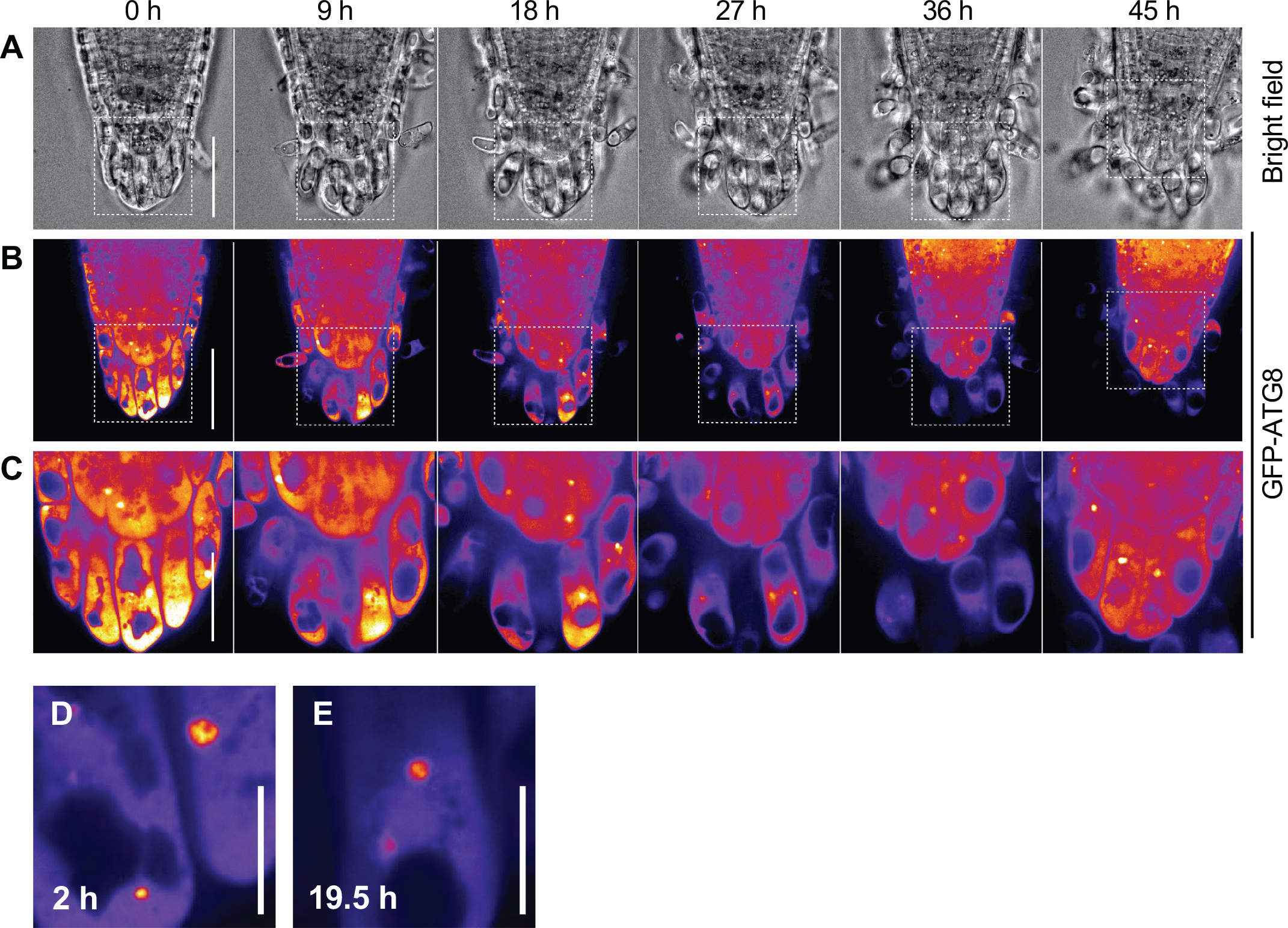
Autophagosomes do not form in the detaching root cap cells of *atg5-1* Time-lapse images of the *35Spro:GFP-ATG8a atg5-1* root tip. Bright-field (A) and GFP- ATG8a fluorescence images (B, C) are shown. Images in (C) are magnified views of boxed regions in (B) of respective time points. Note that the GFP-ATG8a signals were uniformly distributed throughout the cytosol. Occasionally observed punctate signals did not form a donut-shape typical of an autophagosome (D, E). Elapsed time after the start of observation is indicated at the top. Scale bar, 50 µm (A, B); 20 µm (C); 10 µm (D, E). A corresponding video is available as Supplementary Movie S5.

**Fig. S5.**
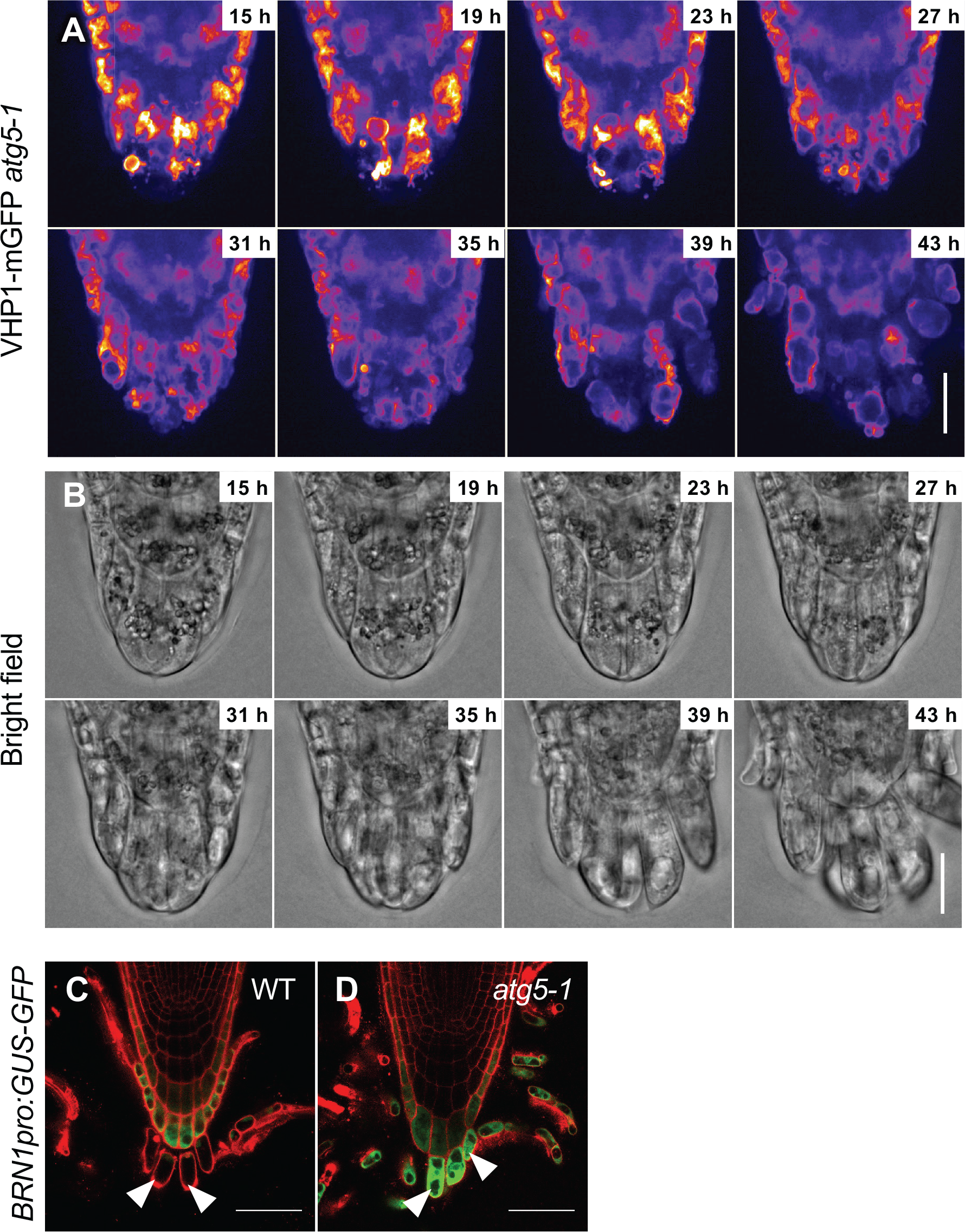
Vacuolization and cytosol digestion do not occur in detaching *atg5-1* cells **(A, B)** Time-lapse images showing vacuolar morphology by the tonoplast-localized VHP1-mGFP fluorescence (A), and corresponding bright-field images (B) in *atg5-1*. In the outermost cells, vacuoles are initially small and fragmented and gradually expand as those in wild type, but fail to expand fully (43 h). Elapsed time after the start of observation is indicated at the upper right corner of each panel. Corresponding video is available as Supplementary Movie S6. **(C, D)** Cytosolic GUS-GFP proteins expressed under the outer layer-specific *BRN1* promoter revealed cytosol digestion in the detaching root cap cells of wild type, as compared with its retention in *atg5-1* (white arrowheads). Scale bar, 20 µm (A, B); 50 µm (C, D).

**Supplementary Movie S1.**
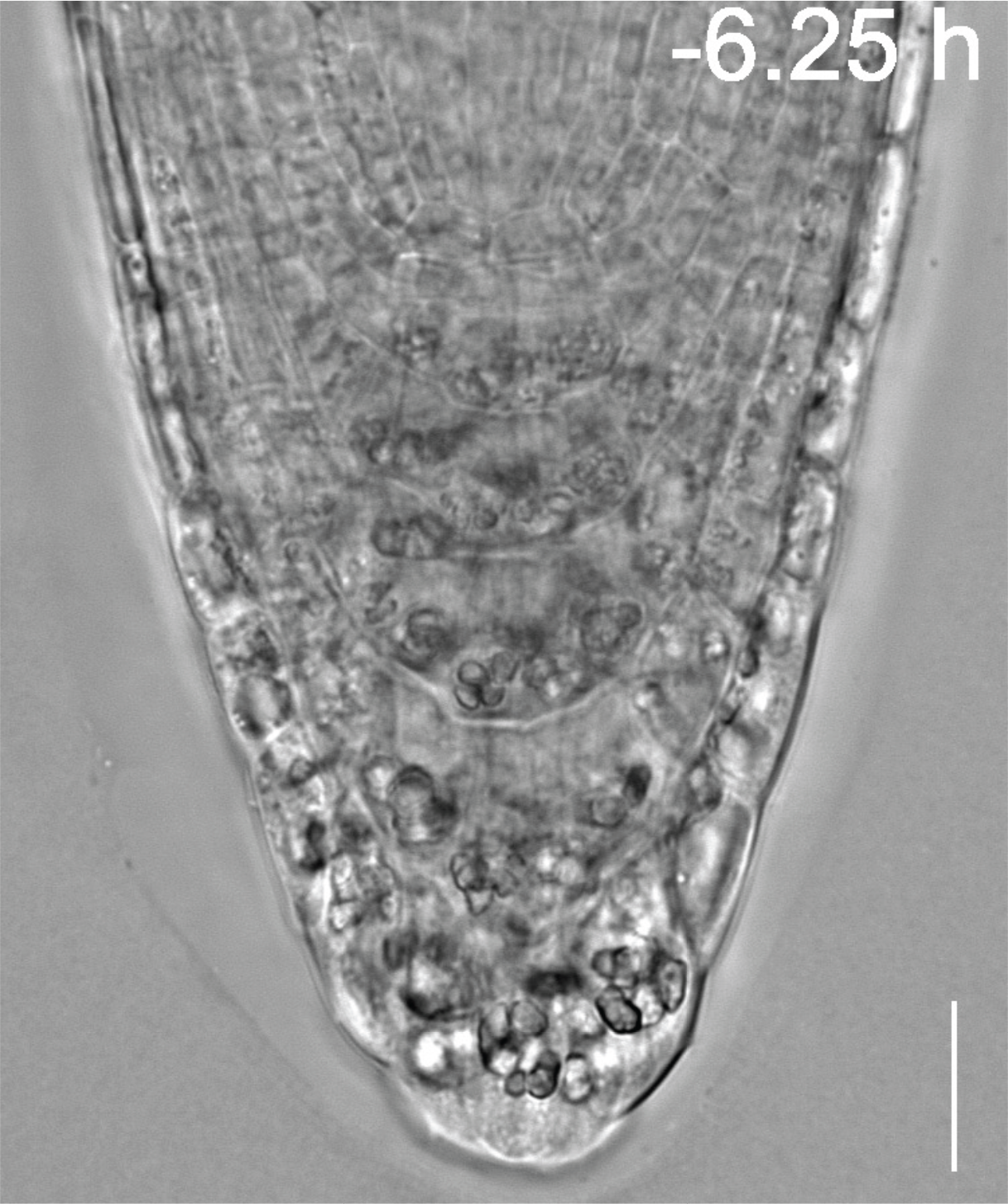
Time-lapse movie showing root cap cell detachment and organelle rearrangement in wild-type root cap cells Scale bar, 20 µm.

**Supplementary Movie S2.**
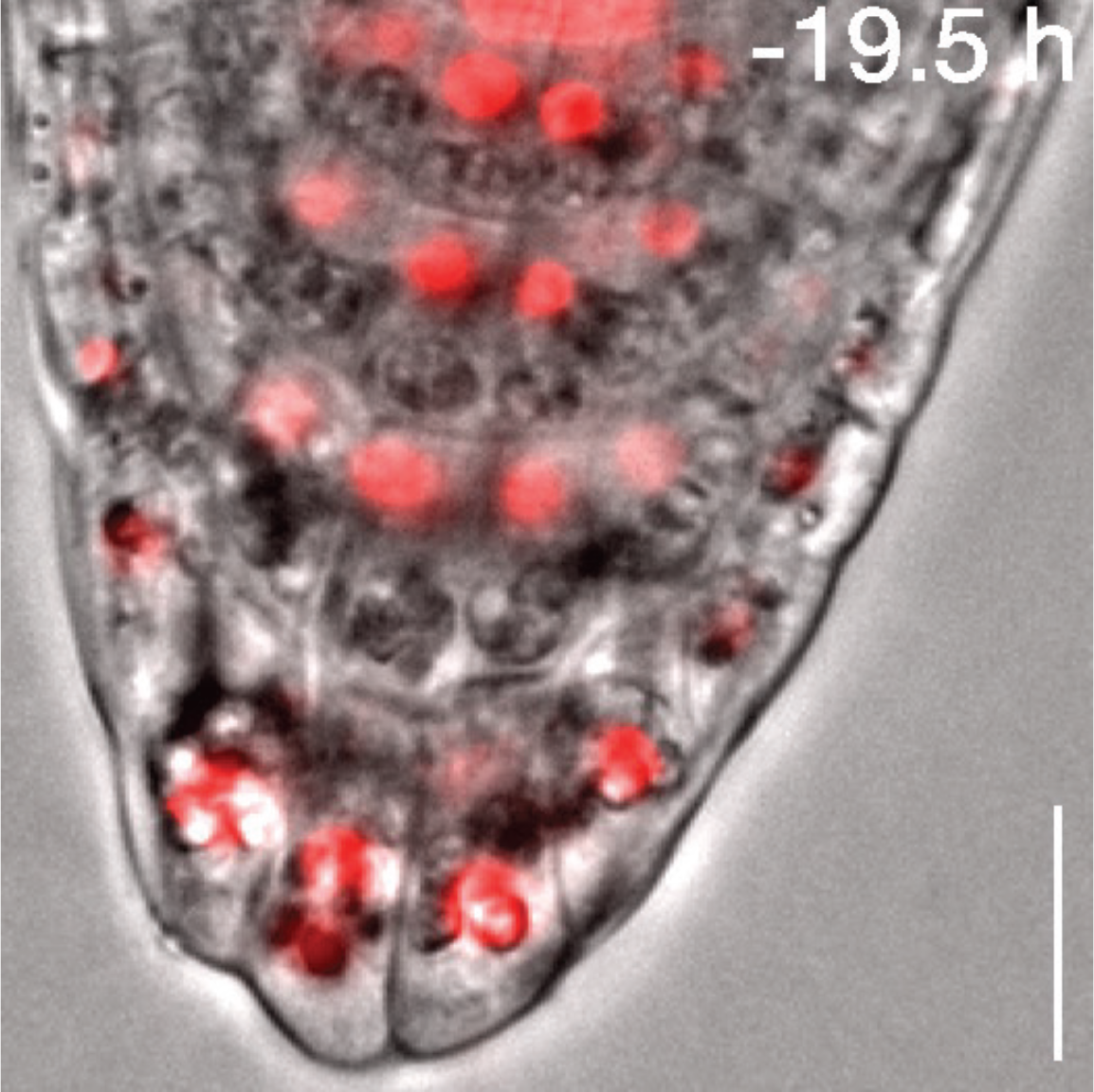
Time-lapse movie showing intracellular relocation of nuclei (red, *DR5v2:H2B-tdTomato*) and amyloplasts (gray particles in the bright field) in the root cap cells Scale bar, 20 µm.

**Supplementary Movie S3.**
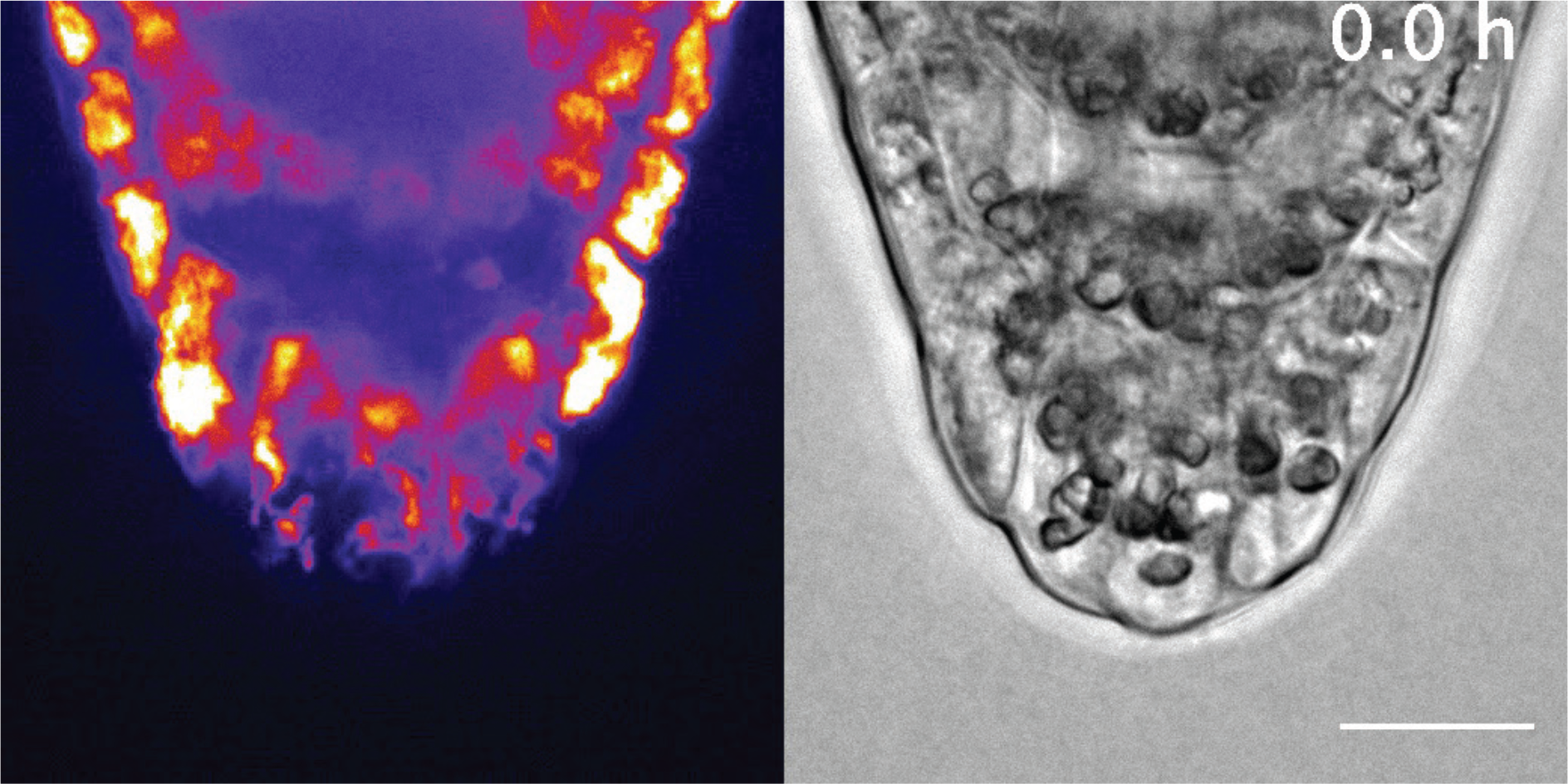
Time-lapse movie showing morphological transition of vacuoles during cell detachment Scale bar, 20 µm.

**Supplementary Movie S4.**
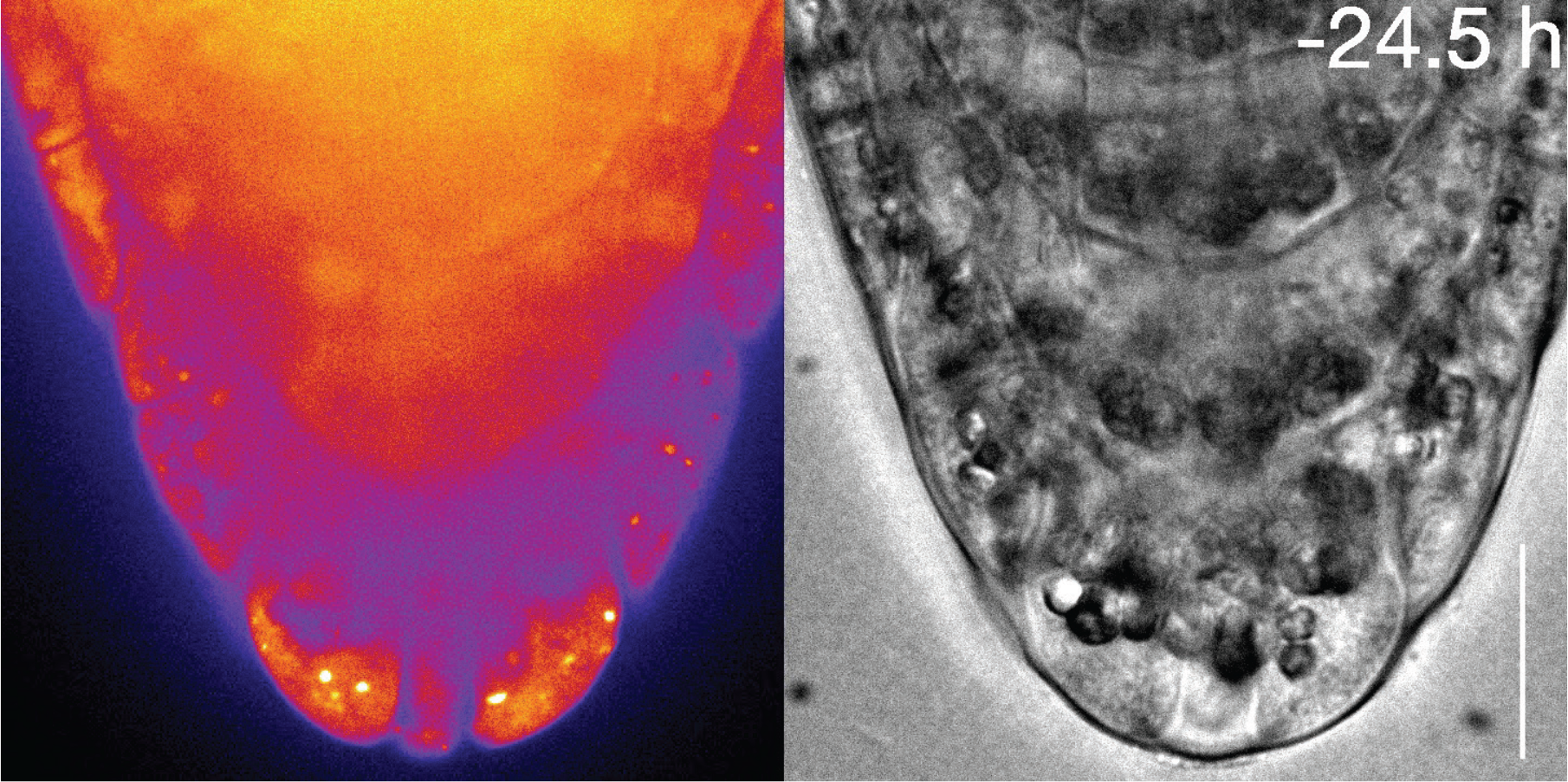
Time-lapse movie showing autophagosome formation in the outermost root cap cells visualized by *35Spro:GFP-ATG8a* Scale bar, 20 µm.

**Supplementary Movie S5.**
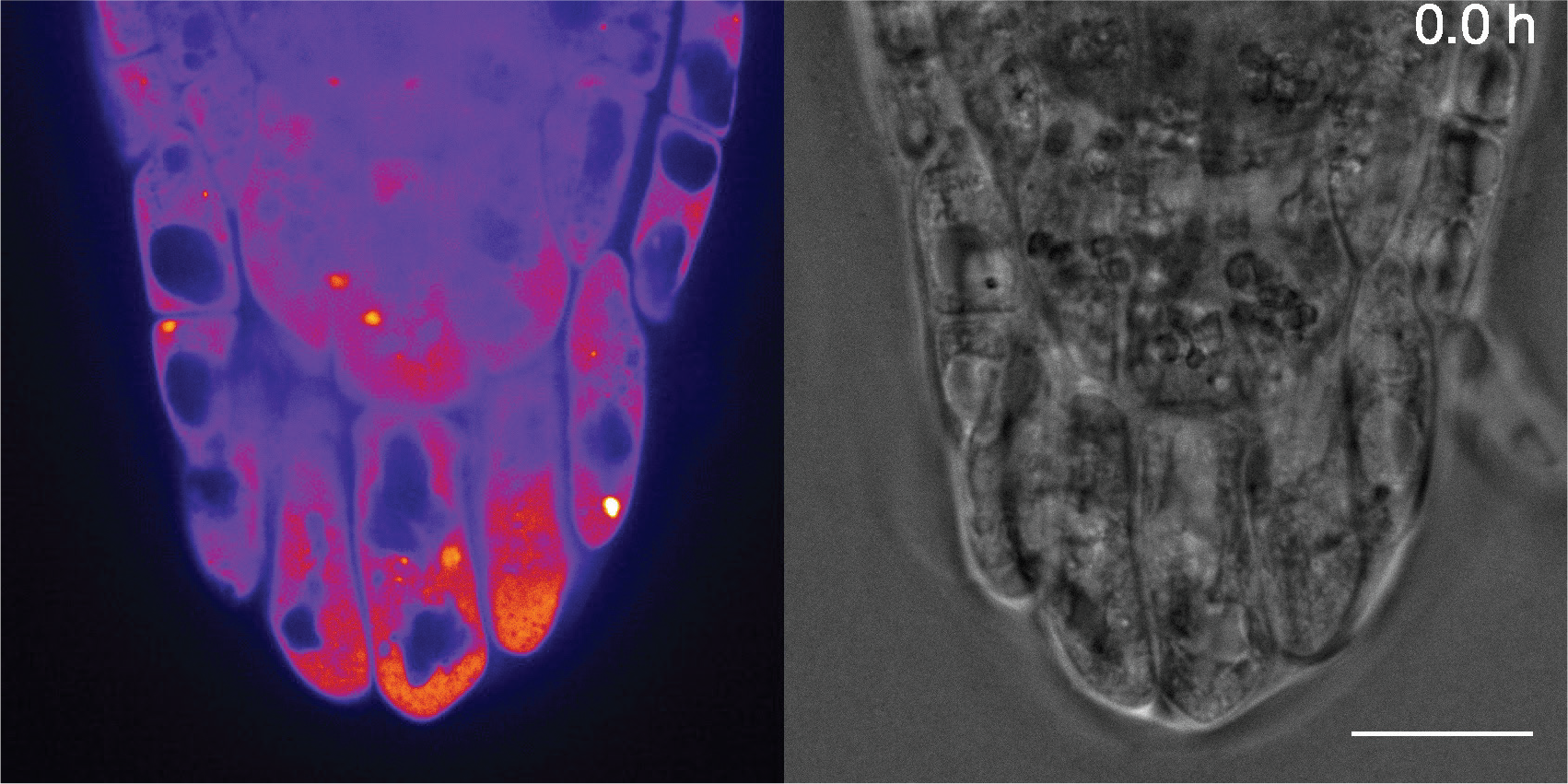
Time-lapse movie showing the absence of autophagosome formation in *35Spro:GFP-ATG8a* in *atg5-1.* Scale bar, 20 µm.

**Supplementary Movie S6.**
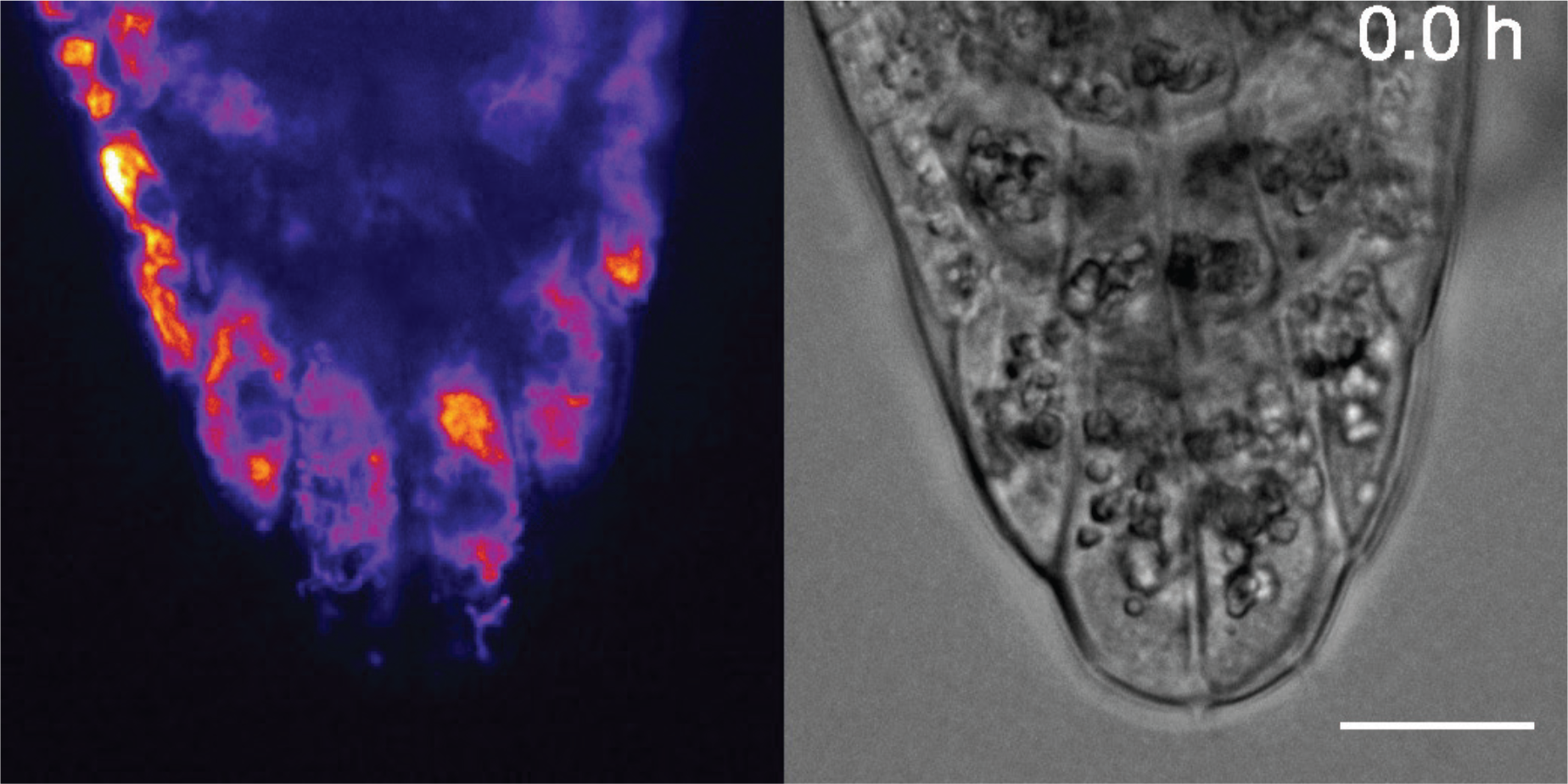
Time-lapse movie showing morphological transition of vacuoles during cell detachment in *atg5-1.* Scale bar, 20 µm.

**Supplementary Movie S7.**
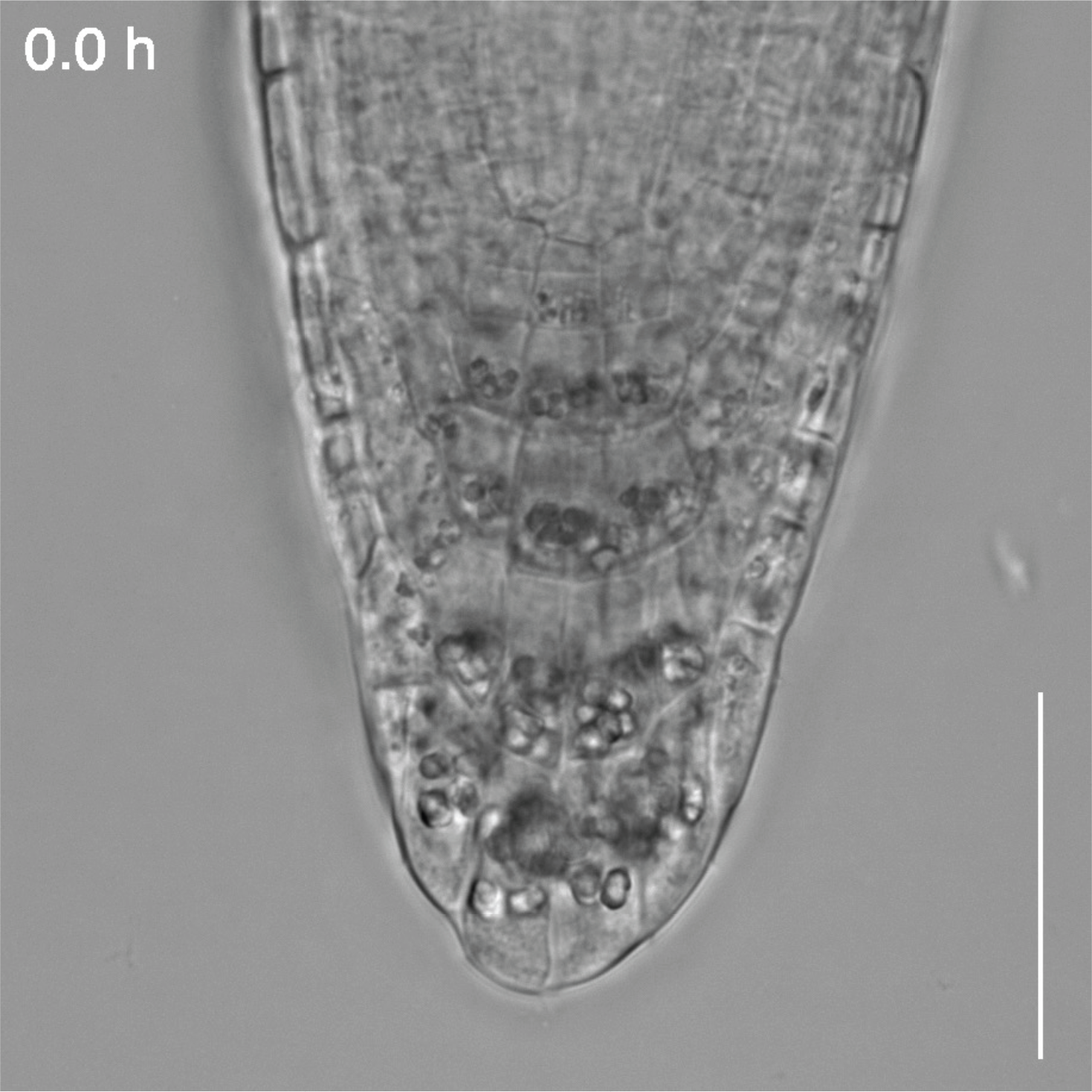
Time-lapse movie showing root cap cell detachment in the wild type Scale bar, 50 µm.

**Supplementary Movie S8.**
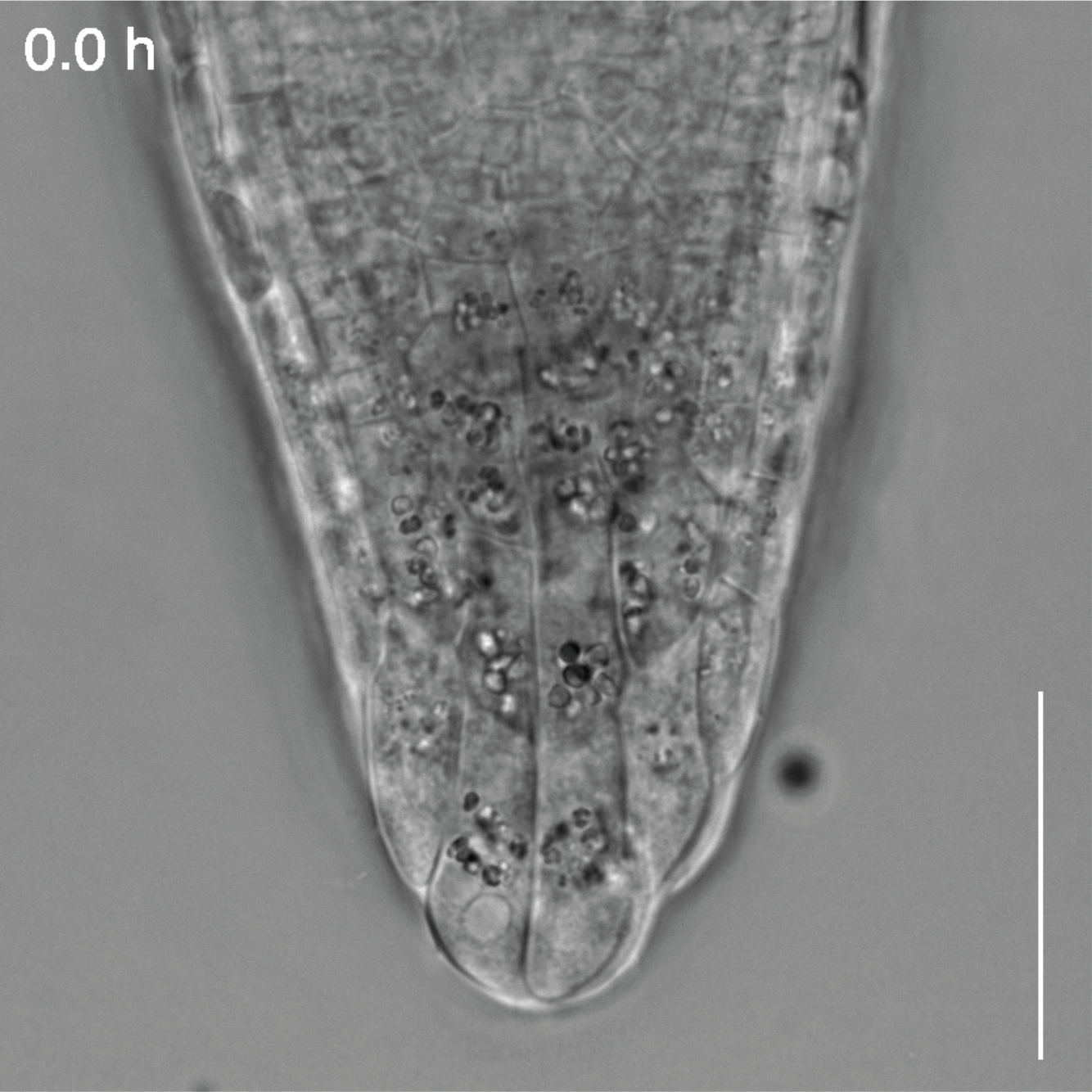
Time-lapse movie showing root cap cell detachment in *atg5-1* Scale bar, 50 µm.

**Supplementary Movie S9.**
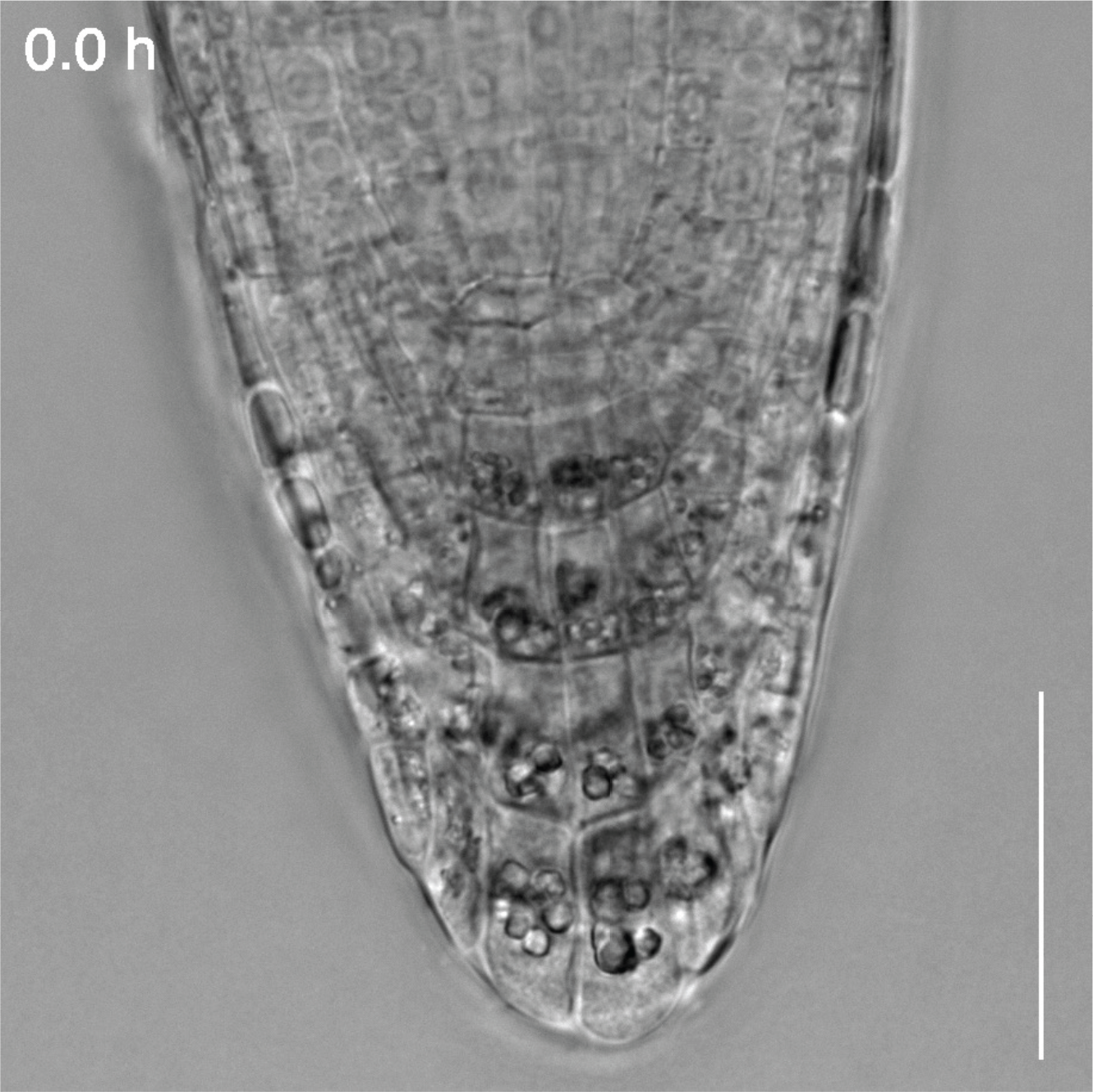
Time-lapse movie showing root cap cell detachment in *atg5-1* complemented with *ATG5pro:ATG-GFP* Scale bar, 50 µm.

**Supplementary Movie S10.**
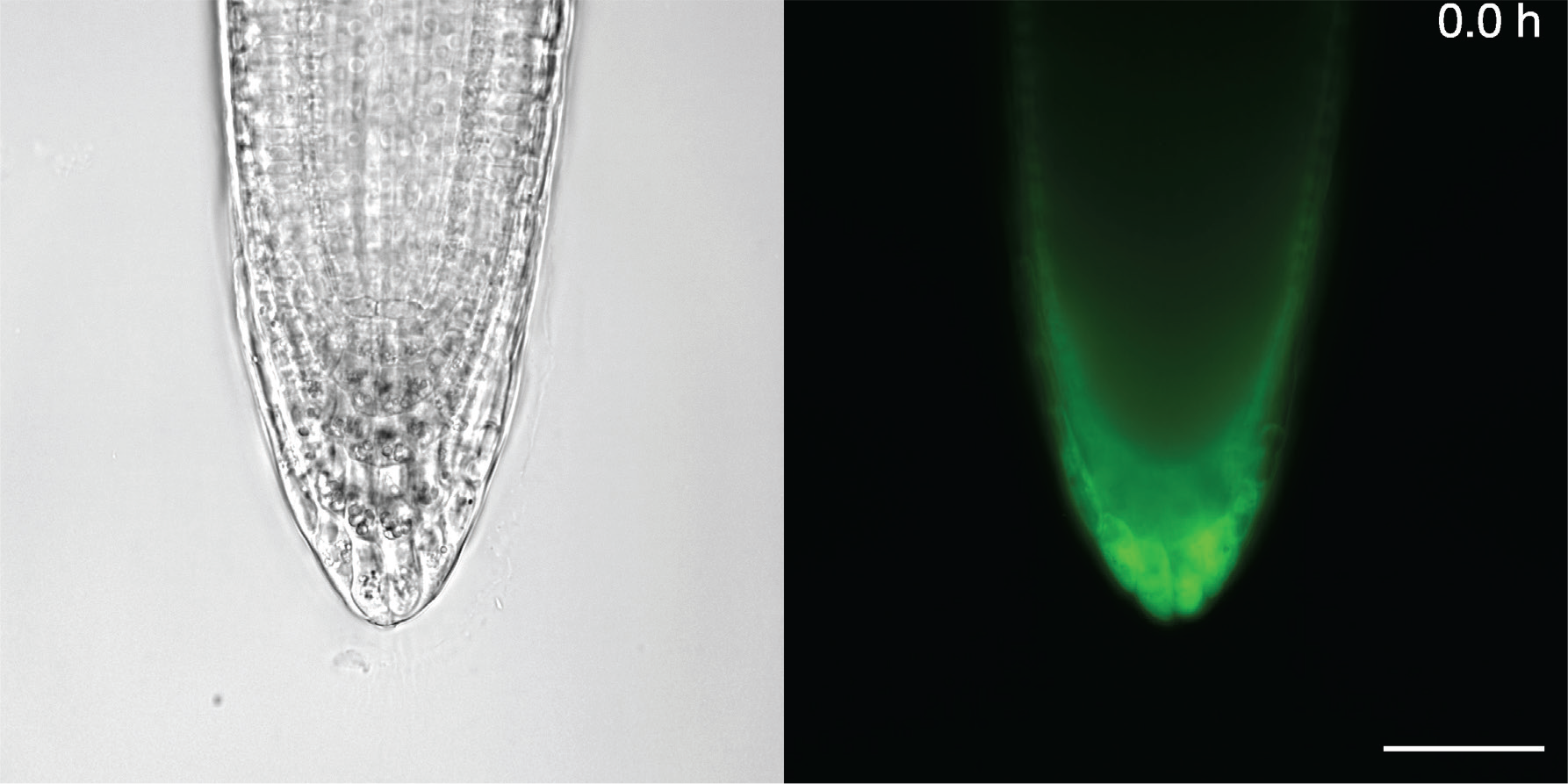
Time-lapse movie showing root cap cell detachment in *atg5-1* complemented with *BRN1pro:ATG-GFP* Scale bar, 50 µm.

**Supplementary Movie S11.**
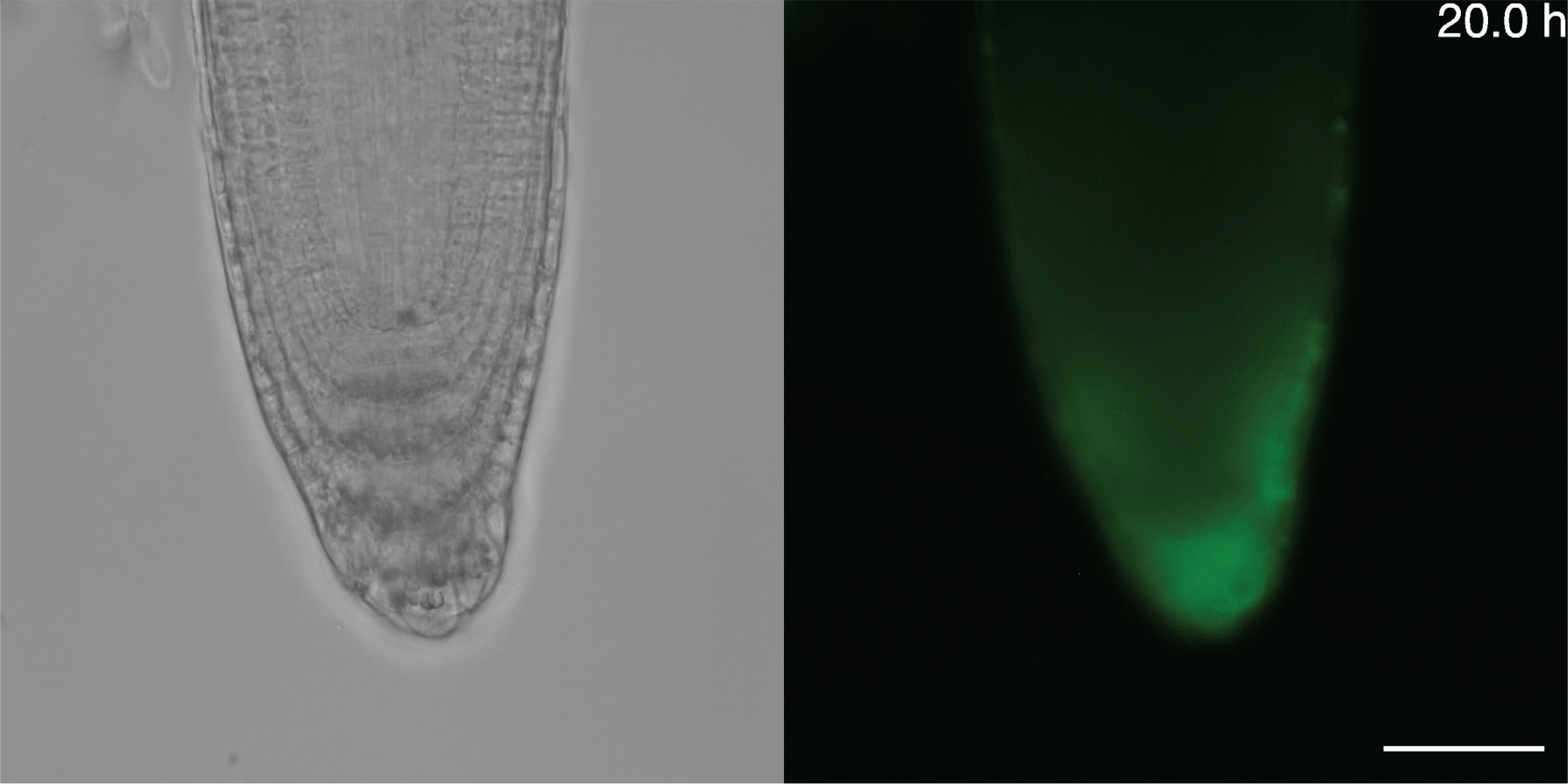
Time-lapse movie showing root cap cell detachment in *atg5-1* complemented with *RCPG1pro:ATG5-GFP* Scale bar, 50 µm.

